# Task-Related Hemodynamic Responses Are Modulated by Reward and Task-engagement

**DOI:** 10.1101/463042

**Authors:** Mariana M. B. Cardoso, Bruss Lima, Yevgeniy B. Sirotin, Aniruddha Das

**Author notes:** To whom all enquiries should be addressed, or: Dept of Neuroscience & Zuckerman Institute, Columbia University, Jerome L. Greene Science Center, Rm L5-005, 3227 Broadway, New York, NY 10027. The work was supported by National Institutes of Health (NIH) Grants R01EY025330, R01EY025673, R01 EY019500 & R01 NS063226 to A.D, and a National Research Service Award to Y.B.S. as well as grants from the Columbia Research Initiatives in Science and Engineering, the Gatsby Initiative in Brain Circuitry, and The Dana Foundation Program in Brain and Immuno Imaging and the Kavli Institute for Brain Science (to A.D.). B.L. received a fellowship from the The Italian Academy for Advanced Studies in America, Columbia University. M.M.B.C was supported by Fundação para a Ciência e a Tecnologia (FCT), scholarship SFRH/BD/33276/2007. Thanks to Maria Bezlepkina and Elena Glushenkova for technical support and lab management, to Liam Paninski for suggestions about analyzing slow changes in physiological responses, and to David Heeger, Mike Shadlen, Elisha Merriam, Charles Burlingham for comments.

## Abstract

Hemodynamic recordings from visual cortex contain powerful endogenous task-related responses that may reflect task-engagement distinct from attention. We tested this hypothesis with hemodynamic measurements (intrinsic-signal optical imaging) from monkey V1, while the animals’ engagement in a periodic fixation task over several hours was varied though reward size, and as animals took breaks. With higher rewards, animals appeared more task-engaged; task-related responses were more temporally precise at the task period (~ 10-20 seconds), and modestly stronger. Surprisingly, 2-5-minute blocks of high-reward trials led to ramp-like decreases in mean local blood volume; these reversed with ramp-like increases during low reward. The blood volume increased even more sharply when the animal shut his eyes and disengaged completely from the task (5-10 minutes). We propose a mechanism that controls vascular tone, likely along with local neural responses, with phasic and tonic components tied to task-engagement.

## Introduction

The use of fMRI (Functional Magnetic Resonance Imaging) in humans, complemented with electrode measurements from animal studies, has considerably advanced our understanding of cortical visual processing. This combination of tools has been particularly useful in understanding exogenous, stimulus-evoked responses. Models of neural responses in humans based on electrophysiological recordings in animals, combined with linear models linking neural to hemodynamic responses, have been effective in accounting for stimulus-evoked fMRI measurements in human subjects and in quantitatively predicting the corresponding sensory percepts^1–9^.

However, fMRI measurements from subjects performing visual tasks also contain large endogenous hemodynamic responses in the absence of, or independent of visual stimuli, even at the earliest stages of visual processing^10–15^. There are at least two types of endogenous response, ‘attention-like’ and ‘task-related’^16^. Unlike the case with exogenous responses, there has been mixed success in interpreting these endogenous hemodynamic responses. Selective visual attention has been characterized extensively through studies in human fMRI^10–15^ with close parallels seen in animal electrophysiology^17–25^. While likely driven by a unified mechanism^26^, attention can take different forms. It could be selective for stimulus location ^10,11,27–29^, or features (e.g. color vs. motion^30^) or timing^28^. The related hemodynamic responses reflect corresponding attributes of the expected stimuli. Attentional responses also increase in strength along the visual cortical hierarchy^11,20^.

Much less is known about the task-related endogenous hemodynamic response, including whether it comprises one or multiple types. It appears to be distinct from selective attention. It entrains to task structure and extends over large sections of cortical areas (e.g. primary visual cortex, i.e. V1) independent of the stimulus^16, 31–33^, where it can even be substantially stronger than stimulus-selective responses^34^. It is also strongest in V1 and progressively weaker in higher visual areas^16^. These difference may reflect distinct brain processes underlying these two endogenous responses. There is growing evidence of the importance of the task-related endogenous response. It may play a role in sensory processing, in temporally grouping otherwise unrelated sensory stimuli^33^ or in switching between stimulus modalities^35^. As yet relatively little is understood about the mechanism of the task-related response even though its presence has been known for over a decade ^16,33,35–41^. This is largely due to the paucity of studies comparing hemodynamics with electrophysiology in alert, behaving subjects.

The current work derives from a task-related hemodynamic response measured using intrinsic-signal optical imaging^42,43^ in V1 of alert macaques performing cued visual tasks^31^. The observed task-related response entrained to task timing independent of visual stimulation, with amplitudes that could compare with or even exceed vigorous visually evoked responses^44^. It appeared to be spatially non-selective, being homogeneous over the optical imaging window and presumably extending beyond^32^. It is thus likely a good model for investigating the mechanism underlying the task-related response seen in humans. Notably, concurrent electrode recordings showed it to be poorly predicted by changes in local firing rates or field potential (LFP) power at any frequency band^31^, unlike stimulus-evoked hemodynamic responses that were well predicted by local electrophysiology^44,45^. Additionally, at a vascular level, this response corresponded to a coordinated contraction-dilation cycle engaging the arterial blood supply into the imaged cortical region^31^. These observations suggested an underlying mechanism distinct from exogenous, stimulus-evoked responses.

Here we explore the link between this task-related hemodynamic response and the level of engagement in a task. The link was suggested by earlier measurements showing correlations between the measured task-related response and task performance^32^, as well as with sympathetic-like markers of mental effort in a task^46^ such as phasic pupil dilation^31^ and heart rate fluctuations^31^. To modulate the level of engagement, we changed reward size systematically^47^ while the monkeys performed a periodic visual fixation task over several hours. Using intrinsic-signal optical imaging and electrophysiology, we looked for effects on the measured task-related hemodynamic response at multiple time scales: of individual trials (^~^10-20 seconds); of blocks of trials (150-300 seconds), and finally, of extended segments of task engagement vs. disengagement as the animal switched between working, as vs. resting with eyes closed (many minutes). Based on our results, we propose that the task related hemodynamic response reflects mechanisms that entrain brain processing more sharply to a task during periods of higher task engagement, possibly as a means of temporally filtering or binding components of a task. Additionally, we propose that the mechanism is accompanied by an overarching pattern of vascular control that enhances vascular tone over both phasic and tonic time scales during a task in coordination with ongoing changes in the level of engagement. Understanding these links would be an important step forward in understanding the dynamic allocation of brain resources in the context of a task.

## Results

### Overview

Two male rhesus macaques performed a cued, periodic visual fixation task, receiving juice reward following every correct fixation with no time out or other punishment for errors (see **Methods**). The task is known to evoke a robust task-related hemodynamic response in the monkeys’ V1 independent of visual stimulation^31,44^. Here we systematically manipulated the size of the reward per correct trial, as a means of modulating the animals’ level of engagement in the task. This was done either in alternating blocks of high and low reward, or in sequences of progressively changing reward (see **Methods**). We recorded V1 hemodynamics using intrinsic-signal optical imaging^42^, a high-resolution optical analog of fMRI^48–50^. This technique deduces brain hemodynamic responses at the exposed cortical surface by measuring changes in reflected light intensity at wavelengths absorbed by hemoglobin. Here we used a wavelength tuned for imaging changes in blood volume (see **Methods**). Imaging was combined with concurrent extracellular electrode recording of multi-unit spiking and LFP. All experimental procedures were performed in accordance with the NIH Guide for the Care and Use of Laboratory Animals and were approved by the Institutional Animal Care and Use Committees (IACUC) of Columbia University and the New York State Psychiatric Institute.

We observed distinct effects on the task-related V1 hemodynamic responses at the three different time scales tested. At the shortest time scale, of individual trials – a few seconds – higher reward led to crisper temporal alignment of the task-related response to each trial, accompanied by a significant if modest improvement in response amplitude. At a slower time scale of blocks of alternating high vs low reward (10 to 20 trials, i.e. 150 to 300 sec per block), we observed consistent alternating ramp-like changes in the mean local cortical blood volume. Notably, the sign of the ramps was such as to decrease blood volume for blocks of high reward while increasing it for low. Finally, periods of disengagement from the task where the animal shut his eyes and rested over many minutes led to further large, sustained increases in the mean local blood volume. None of these effects at any time scale could be accounted for by changes in local spike rate.

The majority of the reported results came from tasks performed in essentially complete darkness (‘Dark-Room Fixation’ *N* = 30 sites, 3 hemispheres, 2 animals). This allowed measurement of the effects on the endogenous task-related hemodynamic response while minimizing exogenous visual confounds^31^. A complementary section (*N* = 33 sites, 2 hemispheres, 2 animals) confirmed that the observed results generalized to the presence of visual stimuli.

### Time scale of single trials: higher reward leads to greater temporal precision

A section of recording made while the animal fixated periodically in the dark illustrates the pattern of task-entrained responses, as well as changes to these responses with reward size (**Fig 1a**). Despite the near total absence of visual stimulation, the V1 hemodynamic recording showed robust task-related fluctuations in local tissue blood volume^31^. These were accompanied, as noted earlier^31^, by phasic sympathetic-like responses^46^ i.e. phasic pupil dilation and heart rate fluctuations, also entrained to the task period. These sympathetic-like responses increased with higher reward. The pupils dilated more per trial, switching dilation size across single trials at block transitions (**Fig 1b**). The mean heart rate fluctuations were stronger (**Fig 1c,e**). Further, animals made fewer errors (fixations broken or never acquired) in high-reward blocks (**Fig 1a, d**). Notably, the mean hemodynamic response also appeared to ramp slowly upwards during the high-reward block, i.e. reducing mean local blood volume, as indicated by the slope of a linear regression line (red, **Fig 1a**); this observation is addressed in a later section on slow changes. Parenthetically we did note a weak fluctuation in recorded spiking that was periodic in the mean and appeared to relate to hemodynamics in some data sets. However, the correlation was unreliable and did not generalize (**Supplementary Fig 1**), consistent with our earlier findings that the task-related response is not predicted by local spiking or LFP^31^.

**Figure 1:**
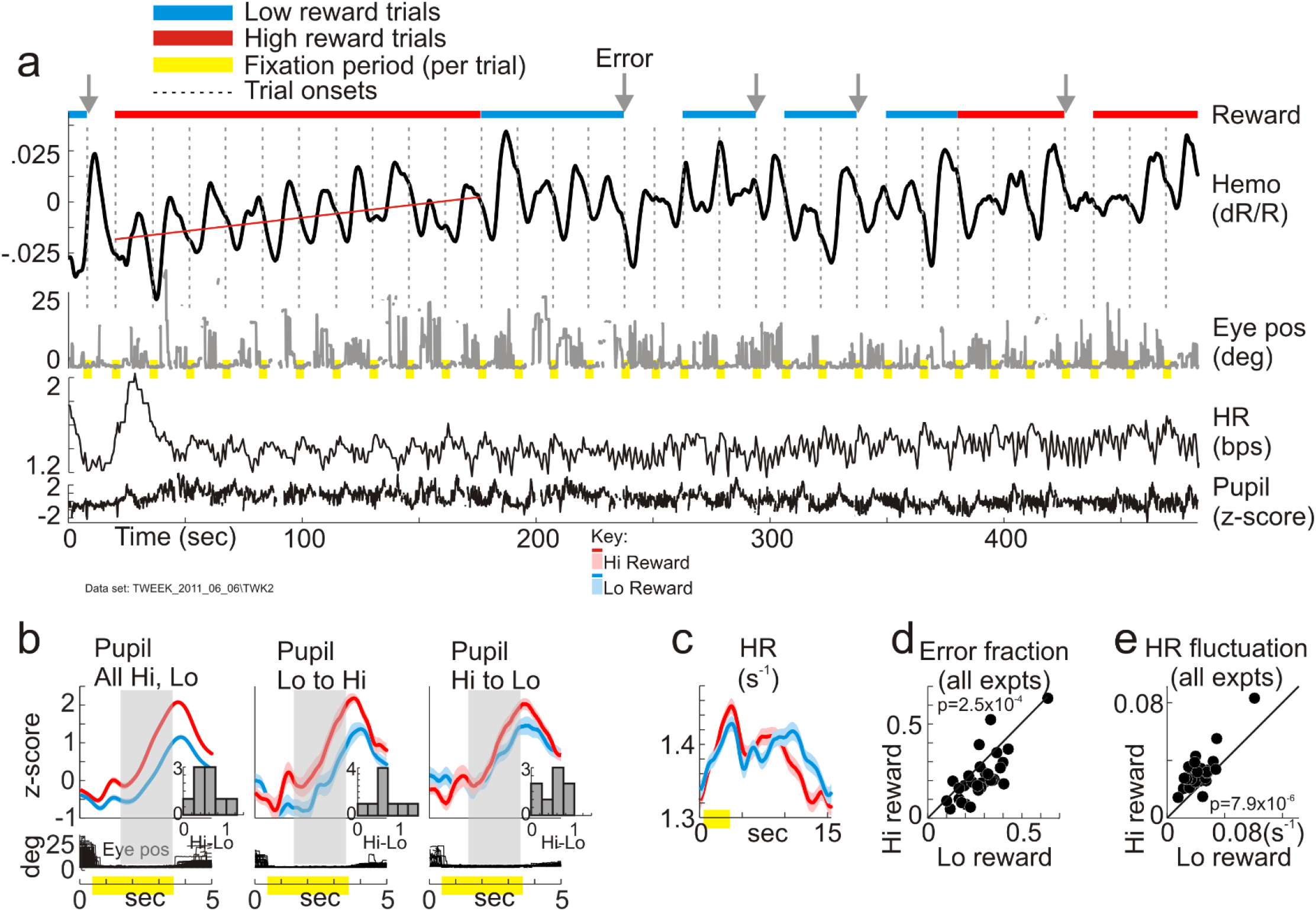
High reward leads to greater engagement in the task**. a-c:** Example dataset, periodic fixation task in the dark. **a:** Continuous records of hemodynamic response (‘Hemo’), radial eye position (‘Eye pos’), heart-rate (‘HR’) and pupil size while reward level alternated between high (0.375 ml per correct trial) and low (0.11 ml) in blocks of 10 correct trials. (Red, High reward. cyan, low. Same color code used all through the paper.) Trials with no color indicate incorrect fixation (compare Eye pos). Each continuous sequence of incorrect trials counts as one error (gray arrows). Monkeys made more frequent errors in low-reward blocks (0.29 for Lo, vs 0.19 for Hi, as fraction of correct trials (N=330)). Hemodynamic response 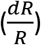 plots fractional change in light reflected off cortical surface; down indicates increasing light absorption, i.e. increasing local blood volume. **b:** Comparing phasic pupil dilation, high vs. low reward trials. ‘All Hi, Lo’: compares all correct high-reward trials (N=160) with low (N=170). ‘Lo to Hi’, ‘Hi to Lo’: compare the first trial following corresponding reward block transitions to the immediately preceding trial linked to opposite reward size (N=15 Lo to Hi transitions, 16 Hi to Lo). Gray shaded rectangle, steady fixation starting one second after fix onset; used for quantifying pupil dilation. Inset histograms show dilation difference (Hi minus Lo reward) for all experiments with reliable pupil recording (N=9). **c:** Comparing amplitude (sd) of mean trial-linked heart rate fluctuations, high-vs. low-reward. (0.038 s^-1^: high, 0.025 s^-1^: low,). Traces in b, c shown as mean +/- SEM (lighter ribbon). **d:** Scatter plot comparing errors as fraction of correct trials, high vs. low reward, all experiments (N=30). **e:** Comparing amplitudes of mean HR fluctuation (sd as in panel c), high vs. low-reward, all experiments. p values in panels d, e: Wilcoxon signed rank test.

At the time scale of individual trials, the primary correlate of high reward on the task-related response appeared to be greater temporal precision, i.e. tighter alignment to trial timing. This was evident qualitatively in lower trial-to-trial temporal jitter for high reward (**Fig 2a,** left panel). The mean of these trial-by-trial responses, averaged across all correct trials, was also higher for high-reward trials. But it was unclear how much of that was due to a true difference in amplitude, as opposed to better temporal alignment of individual responses. To resolve this issue, it was necessary to separately estimate the timing and amplitude of the task-related response for each trial.

**Figure 2:**
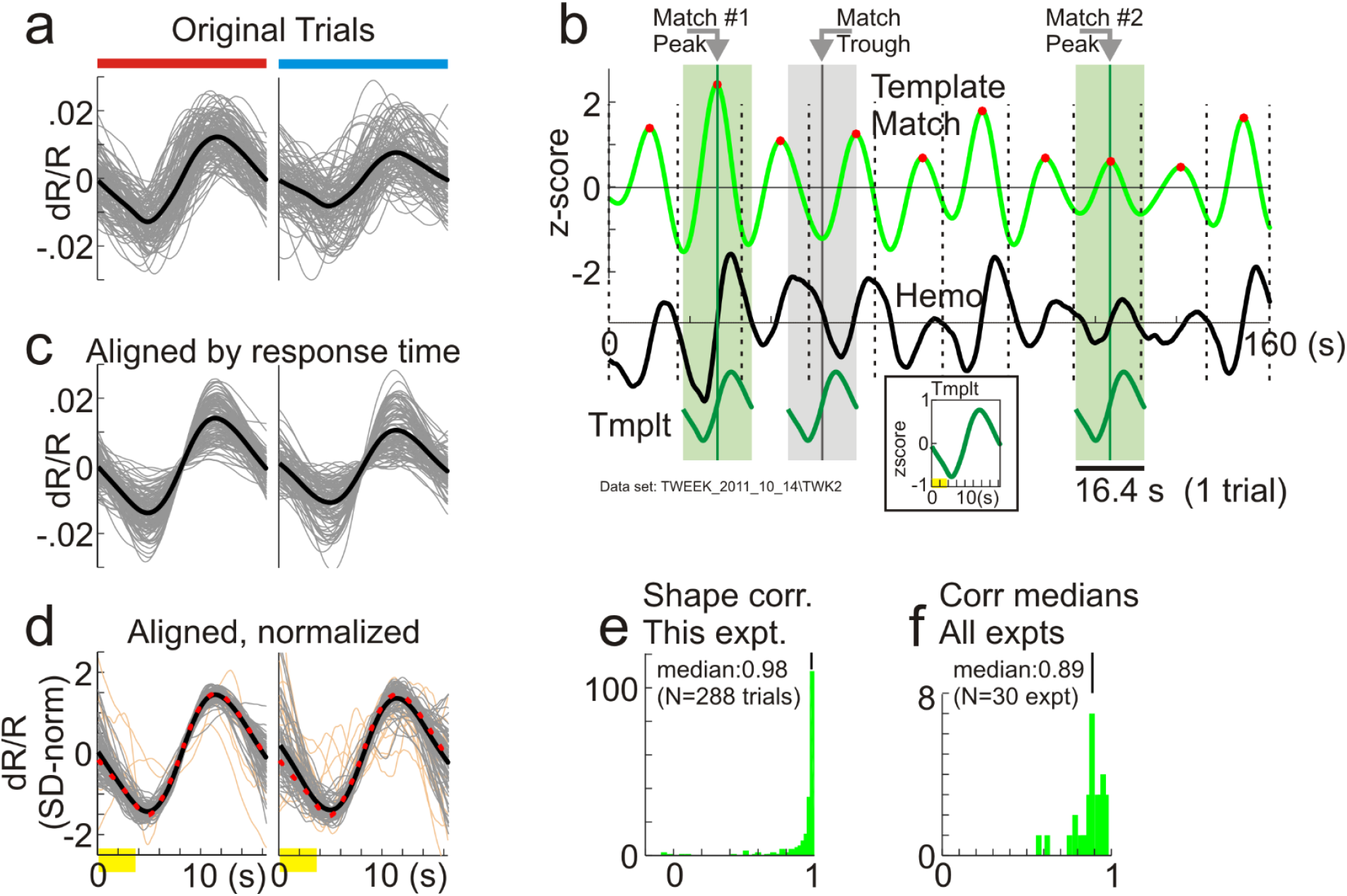
Estimating trial-by-trial timing and amplitude of task-related responses by template matching. **a:** All correct trials in one data set, separated by reward size: high (N=140 trials; left, red bar on top as in Fig 1a) and low (N=148, right, cyan bar). Gray, individual trials, black, mean. Time axis shared with (c), (d) (0=trial onset; yellow, fixation). **b:** Elements of the template match. Black (‘Hemo’), section of recorded hemodynamic response (z-scored, shifted down for visibility); vertical dashed lines, trial onsets. Green (‘Template Match’), sliding-window dot product of ‘Hemo’ with ‘Tmplt’ (inset: defined as mean hemodynamic response, all correct trials). The locations and heights of template match peaks (red dots) define estimated timing and amplitude of task-related response per trial. ‘Match Peak #1, #2’, examples illustrating information carried by peaks. Both #1 and #2 mark locations where Hemo matches Tmplt in shape (see hemo segments on green shading. Compare with ‘Match Trough’, gray shading, phase-reversed Hemo). Greater height of peak #1 vs #2 quantifies higher amplitude of Hemo fluctuation at #1. However, location of peak #2 is better centered in its trial. **c:** Same traces as in (a), aligned by response times estimated from template match. **d:** Same data as (c), normalized by amplitude (sd). Orange, responses with sd in the lowest 10^th^ percentile over the full set. Gray, upper 90^th^ percentile. Black, mean of gray traces. Red dotted line, template. Gray traces match each other and the template well, particularly near midpoint of trial. **e:** Histogram of correlations of aligned responses with the template (Peason’s r; all correct trials, high and low reward, including lowest 10^th^ percentile sd). **f:** Histogram of correlation medians as in (e), all experiments.

We used a template matching approach based on the observation that, other than temporal jitter, Individual responses appeared similar to each other in shape independent of reward size (**Fig 2a**; also see ref^32^). The full hemodynamic recording was thus modeled implicitly as a sequence of task-related responses of stereotyped shape, one per trial, varying only in amplitude and timing from trial to trial. The template was defined to be the trial-triggered average response over all correct trials. This template was slid in a one-trial-long moving window over the recorded response, calculating the normalized local dot product at each time point (‘Template Match’, **Fig 2b, Methods, Eq 1-3**). The dot product is closely analogous to Pearson’s correlation (see **Methods**). We thus surmised that it would have maxima (peaks) at points of high correlation where the recorded hemodynamics locally matched the template in shape– in effect, defining locations of putative task-related responses. But in addition, unlike Pearson’s r which is scale-invariant, dot products scale linearly with the amplitude of their arguments and thus provide a measure of response strength (**Fig 2b**. Also **Methods**). We therefore defined our estimates of task-related response time and amplitude, per trial, to be the location and height of the corresponding template match peak.

After estimating response times and amplitudes as described above, we wanted to check our starting assumption that the measured hemodynamics are well modeled as a sequence of jittered but stereotyped shapes. If the assumption is valid the segments of recorded hemodynamics centered on each peak of the Template Match should match each other closely in shape. To test, we centered each putative task-related response, as picked out through template matching, by its response time as estimated from the same template match (**Fig 2c**). Indeed the realigned responses were strikingly well correlated with each other. This can be appreciated visually by normalizing realigned responses by their amplitudes to help compare shapes (**Fig 2d)**; and quantitatively by correlating realigned responses to the template used for matching (**Fig 2e,f**). The strength of this correlation supports our approach.

With the task-related response times and amplitudes thus quantified we confirmed that the primary effect of higher reward was greater temporal precision. Response times were better aligned to the task period, with consistently tighter distributions (quantified by the 2σ width of the distribution). This was evident for the example data set (**Fig 3a**) as well as in essentially every other data set (**Fig 3c**). High reward also led to significantly higher response amplitude for the example data set (**Fig 3b**). However, that pattern was less consistent over the full set of experiments with only a relatively modest improvement in median response amplitudes overall (**Fig 3d**).

**Figure 3:**
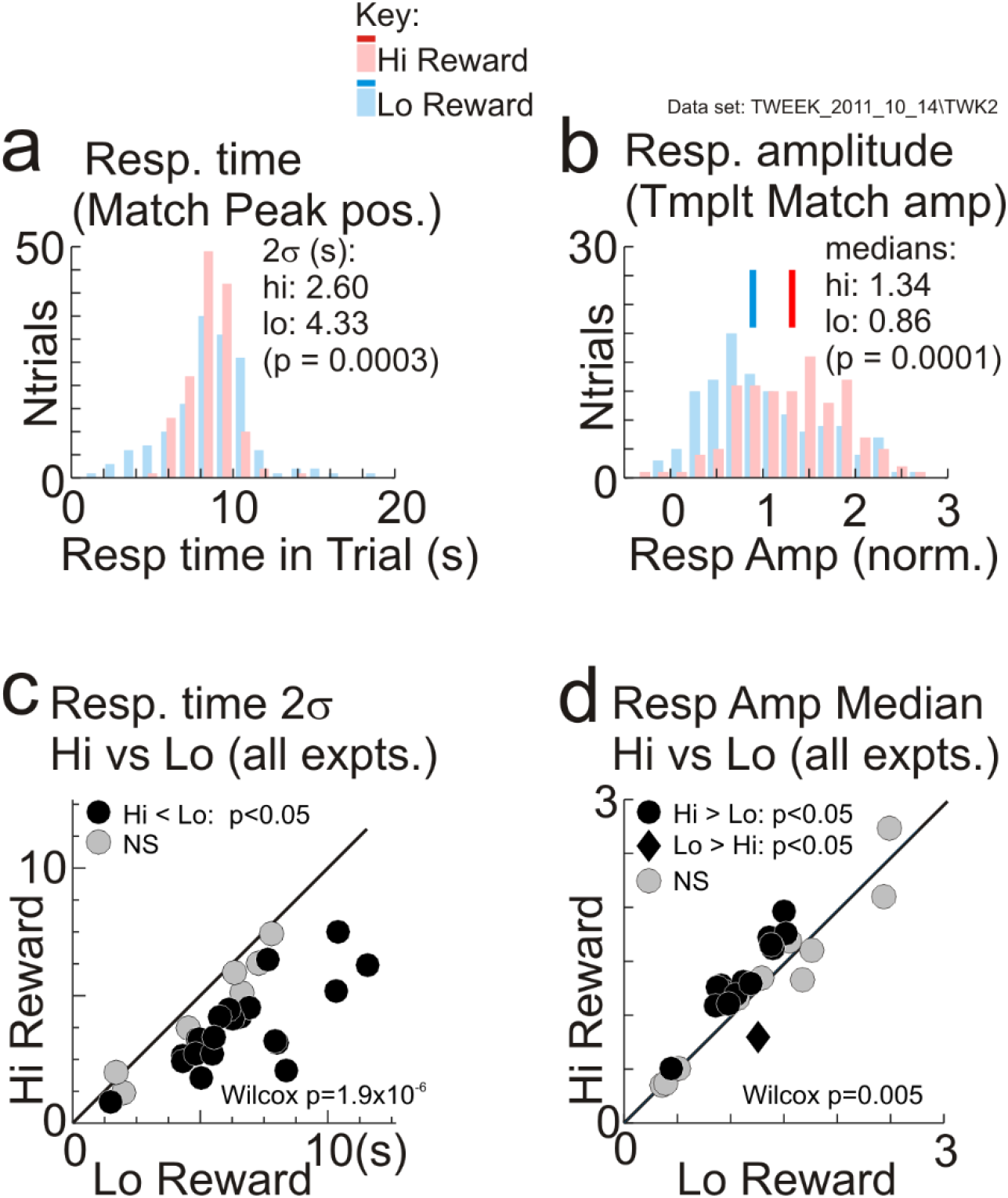
Higher reward leads to more temporally precise task-related responses. **a,b:** Distributions of response times and amplitudes per trial as estimated from template matching, same dataset as in Figs 2a-e. Separated by reward size, with color coding as indicated in key common to panels a, b and to all later figures. Clustering of response times per reward size (a) was quantified as the 2σ width of timing distributions. Response amplitude per reward size (b) was quantified as the median amplitude. Significance (p values) obtained from bootstrap with 10000 resamples. (Median values reported in these and other panels are not the sample medians from the measured distributions but rather, medians from the same bootstrap procedure used to get p values.) **c:** Comparing 2σ widths of response time distributions for high-vs. low-reward trials, per experiment (N=30). **d:** Comparing median response amplitudes for high-vs. low-reward trials, per experiment. p values in (c), (d), Wilcoxon signed rank tests for pairwise comparisons.

We wondered if these results were due to our particular choice of template. We tested by repeating the analysis shown here across all datasets using a range of alternate templates. The alternate templates were also each one-trial long and constructed from measured responses, but using different criteria: for example, phase shifted in time; or using only ‘high signal-to-noise’ responses with amplitudes exceeding a threshold. The task-related-response times and amplitudes estimated by matching to these alternate templates showed a strikingly similar overall relationship to reward size as in Fig 3. This is illustrated in **Supplementary Fig 2** for a particular alternate template with timing and shape distinct from the one used in the main text. This result highlights the overall robustness of our findings. It also suggests that high reward leads to a state of greater temporal regularity and periodicity overall for the duration of the block, accounting for the higher temporal precision in estimated response times independent of the details of the template used.

### Timing precision is robust to noise in template match

We were concerned that the apparently lower temporal precision with low reward could be an artifact of a noisier template match. When task-related responses had lower amplitudes, the template match could be poorer simply due to lower signal to noise. This could lead to noisier estimates of response time with wider distribution and thus apparently poorer precision, but due to the poorer signal to noise alone, independent of reward size (**Fig 4a.** Also consider, e.g., the responses with poor shape match in Fig 2d). Since lower rewards were associated with somewhat lower response amplitudes, this increased noise could make the low-reward responses appear artifactually less precise.

**Figure 4:**
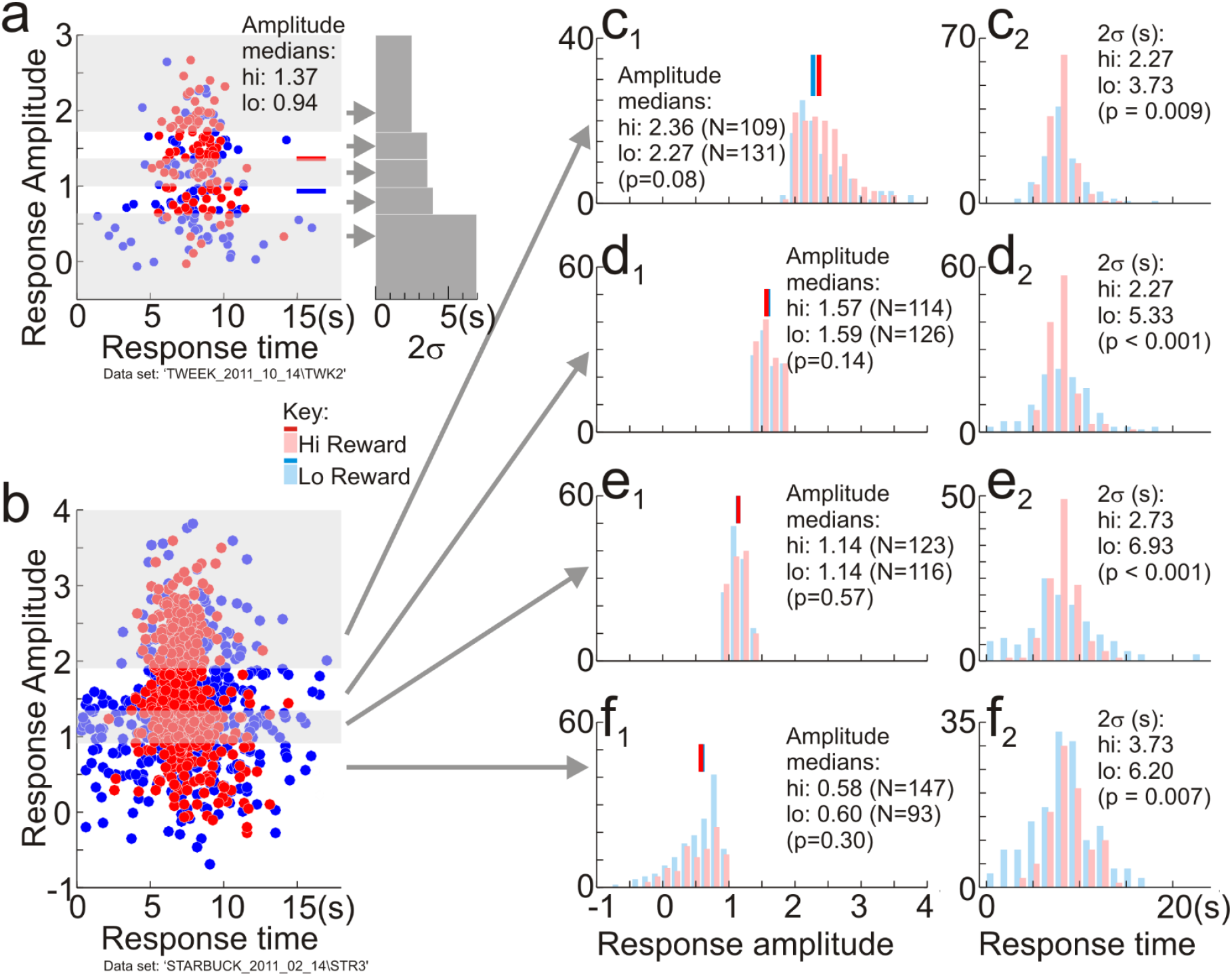
Temporal precision is not an artifact of higher signal for high-reward responses. **a**: An outline of the null hypothesis using a dataset where low-reward responses had substantially lower amplitudes. Left panel. Scatter plot of response amplitude vs. time per trial, colored by reward size. Gray and white shading: quintiles along amplitude axis, combining high and low rewards. Right panel. 2σ width of response time distribution in each quintile. Time axis scaled to match that for left panel. These 2σ widths increased progressively for lower response amplitudes which were also more dominated by low-reward trials. The null hypothesis is that this covariance alone gives low-reward trials larger timing scatter. Arrows: y-axis locations indicate median amplitudes of quintiles. **b:** Plot of response amplitude vs. timing for a large dataset (1285 correct trials; 629: low reward; 626: high reward) separated into quartiles by response amplitude (gray / white shading). Each quartile also roughly matched for numbers of low-vs. high-reward trials. **c1, c2, d1, d2, e1, e2, f1, f2:** pairs of histograms of response amplitude, and timing per quartile. The numbers ‘N’ in parentheses in panels (c1) - (f1) indicate numbers of high and low reward trials. The high-reward responses are significantly more precise than the corresponding low-reward ones in each quartile despite the similarity in response amplitudes (panels (c2) - (f2)). Panels (c1) - (f1) share a common abscissa scale, as do panels (c2) – (f2). P values, bootstrap, 10,000 resamples

To test, we selected subsets from each experiment where the high- and low-reward responses were matched in amplitude and in numbers of data points (**Fig 4b, c1, d1, e1, f1**). If our concern about signal to noise in the template match were valid, high- and low-reward responses in each amplitude-matched subset should exhibit similar distributions of response times independent of reward size. Instead, even after matching for amplitudes, the high-reward responses remained consistently and significantly more temporally precise (See, particularly, Figs 4d1, d2, e1, e2).

### Timing precision is independent of eye fixation timing and eye movements

We wondered if there were some simple oculomotor explanation or correlate of our observation. We considered two possible scenarios under which this could happen.

We considered the null hypothesis that the timing of the task-related response per trial is determined by fix onset, with a stereotyped response time course and hence constant delay following fixation (**Supplementary Figure 3**). If that were the case, response times should be correlated to fix onset times, with unity slope and constant delay. The higher precision with high reward could reflect a behavioral pattern where the animal is more precise in its fix onsets prior to those trials (**Supplementary Figure 3a**). This null hypothesis turned out not to be the case, and task-related response times were uncorrelated with fixation onset. Parenthetically, both animals’ fixation behavior changed over the many months that we tested them intermittently on this task. Initially, both animals tended to maintain fixation for long periods with very few breaks even during intertrial intervals. This led to extended periods of fixation prior to the start of each trial, or even across multiple trials, without any breaks, but unrelated to the timing of the task-related response or reward size (**Supplementary Figure 3b**). Later, both animals showed a different behavioral pattern, moving their eyes around during inter-trial intervals and re-acquiring fixation shortly before trial onset. This led to a pattern of brief fixation periods prior to each trial (**Supplementary Figure 3c**). Task-related hemodynamic response times remained more precise with high reward, independent of the changing pattern of fixation.

We next considered the possibility that animals may have steadier fixation or smaller eye movements during high-reward blocks, due to generally higher engagement in the task (**Supplementary Figure 4**). We failed to see any consistent patterns. There were no consistent differences in fixational jitter between high– and low-reward trials at the resolution of our measurements (60 Hz, 0.33 deg). There were also no consistent differences in eye movements during the intertrial periods where the animals were free to look around. Notably the animals also changed their patterns of intertrial eye movements over the many months of recording. In earlier sessions, they did move their eyes less during high-reward blocks (**Supplementary Figure 4a1-3**). Later, however the animals adopted a behavioral pattern of greater intertrial eye excursions for high-reward trials (**Supplementary Figure 4b1-3**). However, the task-related responses remained more precise for higher reward trials (smaller 2σ width for task-related response time distributions), Independent of this changing pattern of eye movements.

### Timing precision generalizes to the presence of visual stimulation

The question that remained was whether reward size affected task-related responses only in the unnatural circumstance of visual tasks in the near absence of all visual stimulation; or whether such effects generalized to the presence of visual stimuli. To test, we analyzed data from a separate set of experiments where the animals were passively shown visual stimuli – gratings of different contrasts – while performing the same cued, periodic fixation task. For this we first needed to estimate the task-related response from recorded hemodynamics by estimating and removing stimulus-evoked responses. We did so by modeling the overall measured hemodynamics as a linear sum of the stimulus-evoked and task-related components, which we fitted to get the optimal kernels for the two components^51^ (**Methods, Eq 4-6; Supplementary Fig 5**). The optimal HRF kernel thus obtained was then convolved with the recorded spiking to estimate the stimulus-evoked component of hemodynamics and regress it away from the full hemodynamics. The residual – that was, by construction, the component of hemodynamics not predicted by local spiking – was then defined to be the task-related component of the hemodynamic response, equivalent to the full hemodynamic response in the dark-room.

The task-related response thus estimated in the presence of visual stimuli was again temporally more precise with high reward, just as with the task undertaken in the dark room. This can be seen qualitatively after separating the estimated task-related response into individual trials and segregating trials by reward size. These trial-wise responses were visibly less temporally jittered for high reward (**Fig 5a**). The timing and amplitude of these task-related responses were quantified by matching to a template just as for recordings in the dark room; the template was taken to be the optimal mean kernel for the task-related component as estimated from the fit (**Methods, Eq 7,8. Supplementary Fig 5**). The results of this template match closely paralleled those obtained in the dark room fixation task. The estimated response times were again more tightly clustered for high-reward trials, both for this specific dataset (**Fig 5a,c**) and over the set of visually stimulated experiments (**Fig 5e**). Response amplitudes showed only a modest improvement (**Fig 5d,f**). Notably, high-reward trials were also associated with greater pupil dilation (**Fig 5b**).

**Figure 5:**
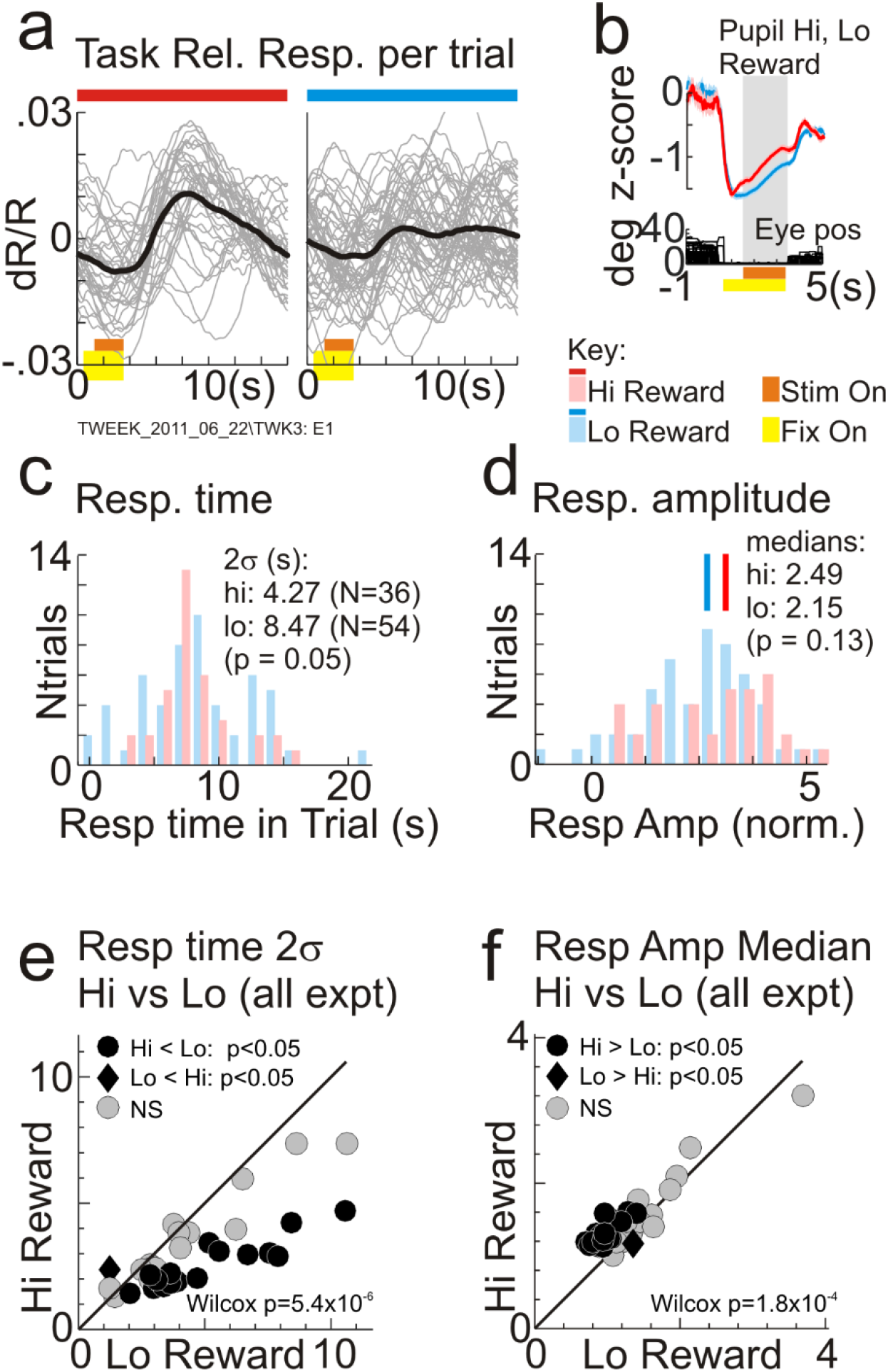
Task-related responses in the presence of visual stimuli are temporally more precise and modestly higher in amplitude for high reward, as in the dark room. **a:** Residual task-related responses separated by trial and by reward size. Obtained by regressing away stimulus-evoked responses (see **Supplementary Figure 5**). **b:** Comparing pupil dilation for high vs. low-reward trials. Gray shading: period over which pupil dilations are compared, starting one second after fixation. (These pupil measurements were made in the presence of visual stimulation unlike dark-room results, Fig 1b; likely accounting for different shape of trace including initial constriction on fixation). **c:** Distribution of response times from template-match, this data set. **d:** Distribution of response amplitudes, this data set. **e:** Pairwise comparison of 2σ widths for high vs. low reward, per experiment (N=33). **f:** Pairwise comparisons of median response amplitudes for high vs. low rewards, per experiment (N=33). p values in (e), (f): Wilcoxon signed rank tests for the pairwise comparisons.

### Time scale of blocks of trials: mean blood volume decreases for high reward, increases for low

Analyses up to this point were restricted to the scale of single trials, i.e. ~10 to 20 sec. However, we also noted slow ramp-like drifts in the mean local blood volume over blocks of ten to twenty trials of a given reward size, i.e. about 150 to 300 seconds (**Fig 1a; Fig 6**). Surprisingly, the ramps decreased blood volume for high-reward blocks while increasing it for low. Regression lines fitted through sequences of correct trials per block clustered into distinct sets of negative slopes (increasing absorption of light during imaging, i.e. increasing blood volume) for low-reward blocks and positive slopes (decreasing blood volume) for high (**Fig 6b, c**).

**Figure 6:**
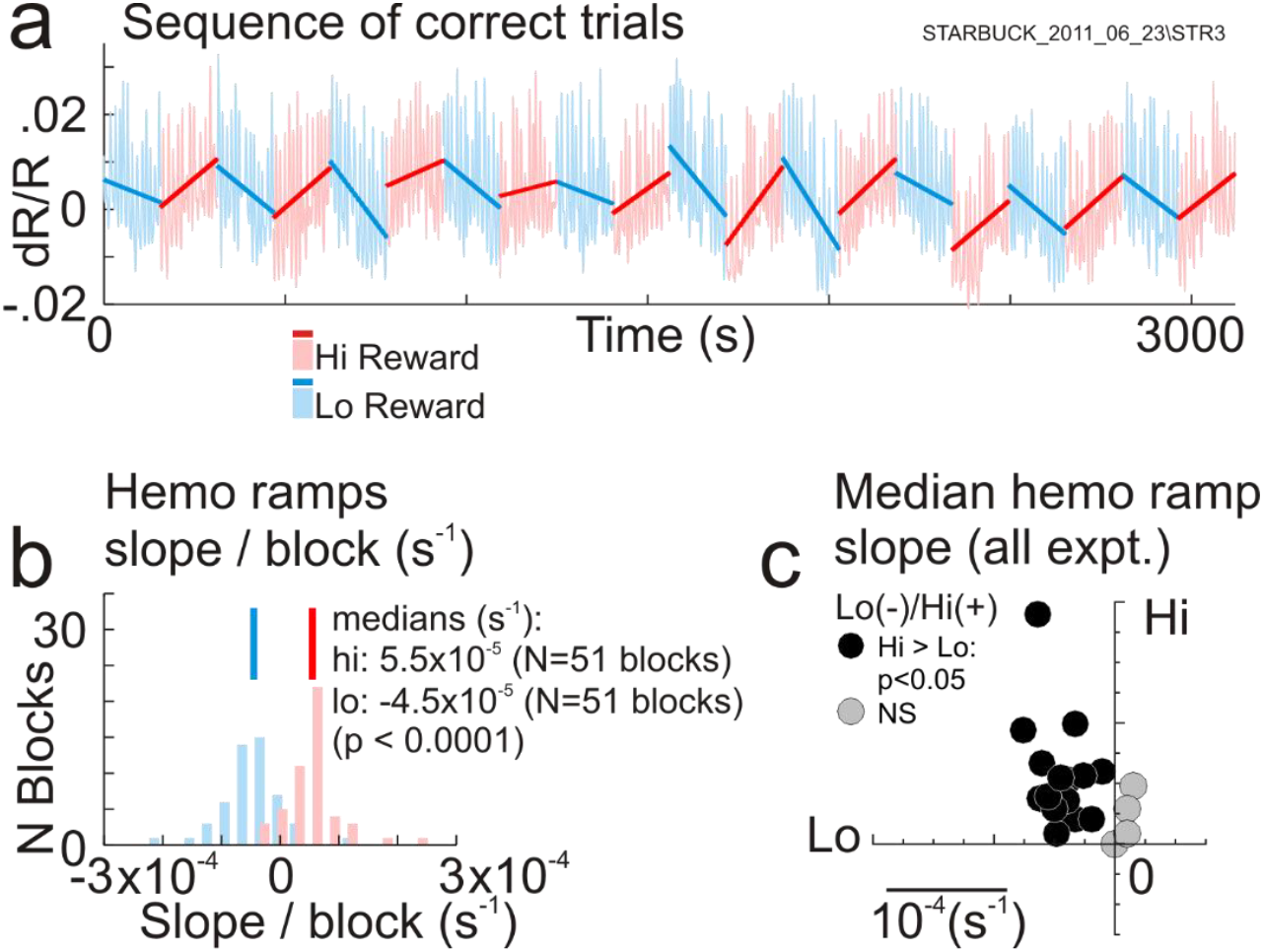
Mean local cortical blood volume increases for low-reward blocks and decreases for high, in alternating ramp-like drifts. **a:** Recordings from a sequence of correct trials, alternating between high and low reward in blocks of 10. Lines indicate regression fits to each block separately. Increasing slope implies decreasing local tissue blood volume. Correct trials were concatenated after excising incorrect ones while maintaining vertical position. **b:** Histogram of regression slopes in high-vs. low-reward blocks. Same data set as (a). (p values, bootstrap, 10000 resamples). **c:** Comparing median slopes of high-reward vs. low-reward blocks over the set of all experiments. All the statistically significant data points lie in the upper left quadrant, Lo(-) / Hi(+), i.e. with negative slopes for low- and positive slope for high-reward blocks (N=19 experiments: using only those with at least 10 pairs of alternating blocks of 10 trials each).

These slow hemodynamic drifts were not driven by slow changes in spiking. Blocks of trials with alternating ramps of mean blood volume failed to show similar alternating ramps of mean spiking (**Supplementary Fig 6**). To test more quantitatively, we first simulated the spiking patterns required to generate the measured hemodynamic slopes on convolving with the corresponding optimal fitted HRF, per experiment (**Supplementary Fig 6 e,f**). The slopes of the simulated spiking ramps alternated in sign with reward size, as expected. Each measured spiking slope was then divided by the slope of its corresponding simulation, to compare. If the measured slopes had the same sign as their simulations, these ratios would consist of positive numbers, with some magnitude reflecting a scale factor. This was not the case; the ratios were equally likely to be positive or negative, for both high and low reward. The measured spiking slopes were thus uncorrelated with those required to generate the measured hemodynamic slopes.

### Switching from alert engagement to rest with eyes closed: further profound increases in blood volume

We wondered if the slow increase in mean local blood volume accompanying reduced reward could be part of a broader pattern of shifts in mean local blood volume accompanying shifts in the level of engagement. A potential clue was seen in the continuous measurements during long dark-room recording sessions lasting up to three hours. In these sessions, in between extended stretches of working well, the animals would take occasional breaks of many minutes where they stopped working and rested with their eyes shut. The mean local blood volume in V1 increased strikingly during these breaks, returning to baseline when the animal resumed working (**Fig 7a**). This pattern appeared to be an extreme manifestation of the ramp-like changes in blood volume with reward size where lower reward, with its lower level of engagement, (**Fig 1**), led to increasing mean blood volume (**Fig 6**).

Before ascribing an association with reduced engagement in the task, we needed to test whether the increased blood volume could be accounted for simply by concurrent changes in neural or physiological drivers. As possible drivers we considered the mean heart rate, and the mean local multi-unit spike rate (recorded separately at two electrodes spaced 4 mm apart in the recording chamber). We also measured the pairwise noise correlation of spike rates between the two electrodes over a one-second sliding window since that was expected to increase at rest^52^. To focus on slow changes all recordings were downsampled to get, in effect, the smoothed average in a 60-sec window (see **Methods**).

We then assessed changes in these physiological and neural measurements as the animal switched state, marking the state based on the fraction of time within the 60-second window that the eyes were closed (Fig **7a**, top row, ‘Eyes closed’). Spontaneous eye closures in the dark have been shown to provide a useful measure of drops in vigilance^53^, correlating well with EEG and fMRI indicators^54^. For this study, epochs where the eyes were shut more than 60% of the time were defined as ‘rest’ while those with less than 5% of eye closure were considered ‘alert’. To check, we compared with an LFP measure of alertness based on the ratio of power in the beta- and theta-range frequency bands (15-25 Hz, and 3-7 Hz respectively) as suggested by earlier studies (ref^54^; also, reviewed in ref^55^). To get a measure that was low when the animal was alert and high when at rest^55^, as with eye closure, we placed the theta power in the numerator. The square root of this ratio further compressed the dynamic range to roughly 0-1 as with eye closure. These two measures, based on eye closure and on the LFP were closely comparable (**Fig 7a**, upper two rows and inset), supporting the use of eye closure to segregate physiological measurements by alertness state.

**Figure 7:**
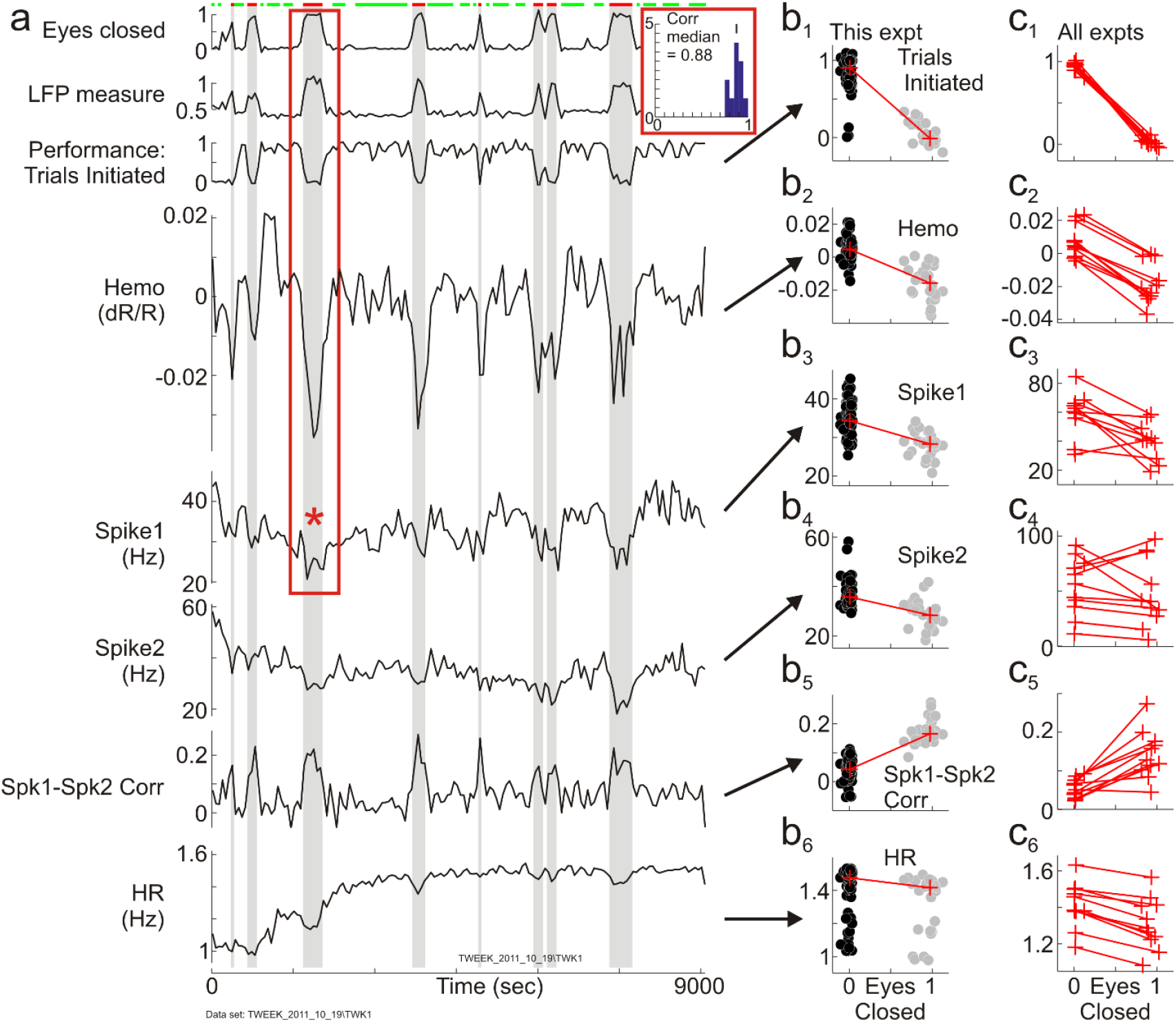
Mean local blood volume, spiking and heart rate track switches between states of alert engagement and rest. Traces show continuous 2.5-hr records of measured variables as indicated by adjoining labels, smoothed and downsampled to a 60-sec sampling rate to track slow changes. (See text and Methods for more details). Red and green bars on top mark epochs of rest (defined as ‘Eyes closed’ > 0.6, i.e. >60% of the 60-sec sample time) and alert engagement (sections with eyes open, defined as ‘Eyes closed’ < 0.05, i.e. <5% of the time). ‘Eyes closed’ is highly correlated with the LFP measure (See text for definition. Pearson’s r = 0.94 for the example dataset. Inset shows histogram of Pearson’s r for similar pairwise correlations over all datasets used for this analysis, N=11). Performance in the task is quantified as the (smoothed) fraction of trials initiated in the 60-sec (i.e. ~ 4-trial) window. The red box indicates the section of hemodynamic and corresponding Spike1 spiking measurements (red asterisk) that are analyzed at a higher temporal resolution in Fig 8a. **b1-b6:** scatter plots of the measured values for the given experiment, as indicated, vs. ‘Eyes closed’. Each data point represents a single smoothed, non-overlapping 60-sec sample. Data points are segregated into ‘alert engaged’ (black) and ‘rest’ (gray) using the value of ‘Eyes closed’ as described. Red lines connect medians. **c1-c6**: lines connecting medians as in panels b1-b6, for all experiments used (N=11).

The mean neural and heart rate measurements, thus segregated, showed systematic changes as the animal switched between states of rest and alert engagement, but in a direction opposite to that expected to increase blood volume at rest (**Fig 7a**). Thus the mean heart rate, averaged over the moving 60-sec window, reduced systematically relative to its local baseline value each time the animal disengaged from work and rested with eyes shut (**Fig 7a: bottom trace (‘HR’)**: see shaded areas indicating rest). This is consistent with the abrupt falls in mean heart rate and blood pressure seen at sleep onset in human subjects^56^. But it suggests that the concurrent increase observed in V1 local blood volume is not a passive consequence of cardiovascular changes as that would require an increase rather than a decrease in heart rate^57^. Similarly, the mean spike rate recorded at individual electrodes typically decreased as the animal rested. If the blood volume at rest were driven linearly by local spiking then the mean spike rate should have increased^1^. Notably, although the mean spike rates at individual electrodes largely decreased, the pairwise correlation of spike rates over the pair of electrodes showed the expected^52^, striking increases for the epochs of rest vs. alert engagement.

Comparing hemodynamics and spiking at the higher, imaging temporal resolution (15 frames/sec) supported our contention that the large blood volume increases at rest are not predicted from spiking. This conclusion was not immediately apparent on qualitative inspection (**Fig 8a**). At this temporal resolution the spiking response showed expected^58^ bursts of high instantaneous spike rate (red arrow, **Fig 8b**) that stood out despite the overall reduction in mean spike rate as the animal rested with eyes shut (see caption, **Fig 8a)**. The corresponding blood volume measurements showed large swings in amplitude that appeared, qualitatively, to follow the bursts of spiking. Our earlier work showed that the recorded hemodynamics is poorly predicted by spiking when the animal is engaged in his task, due to the presence of the task-related response (**Supplementary Fig 1**, refs^31,44,51^). But there should be no task-related response, by definition, when the animal is disengaged from the task with his eyes shut and the hemodynamics could in principle be predictable from spiking. It was thus important to test the relationship between the two at this higher temporal resolution.

**Figure 8.**
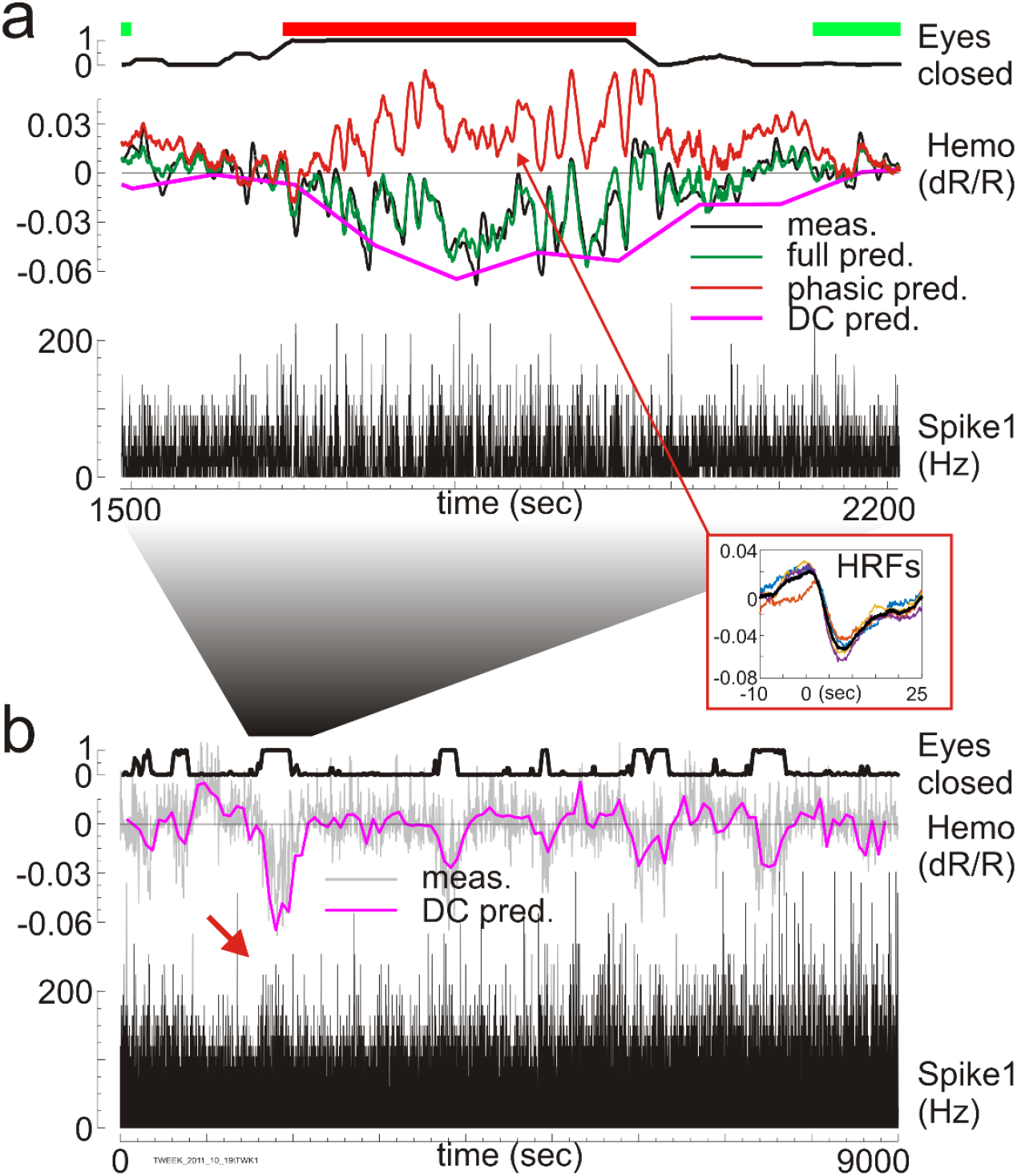
Hemodynamic response during rest is the sum of a component predicted by spiking plus an additional DC shift**. (**Same dataset as in Fig 7, but at a temporal resolution of 66.7 msec (15 Hz camera frame rate).) **a:** Expanded view of the Eyes closed, Hemo and Spke1 traces from section of data enclosed by red box in Fig 7a (also see position relative to full time line, (b)). Green and red bars, ‘Eyes open’ and ‘Eyes closed’ as in Fig 7a, top trace. Bursts in spiking during rest are evident as peaks of high instantaneous spike rate (red arrow, Panel (b)) despite lower mean spike rate over this epoch (24.7 spk/sec average under ‘Eyes closed’ red bar, Panel (a); vs 29.4 spk/sec average in the two flanking green sections where eyes were open. Same data as in Fig 7a, red asterisk). ‘Hemo’ Black, measured. Green, full prediction using optimal deconvolved HRF kernels. Red, prediction using phasic portion of HRF alone. Inset ‘HRFs’: phasic parts of kernels from deconvolution windows in ‘Eyes closed’ segment; color arbitrary (Matlab default). Maroon line (‘DC pred’) shows additional DC component of each deconvolved kernel. **b:** Results of the same deconvolution and prediction shown over the full experiment. Only measured spiking, measured hemo trace and DC prediction are shown; full and phasic predictions are not shown to avoid clutter. Red arrow: burst of high instantaneous spiking despite lower overall mean.

We tested using deconvolution, i.e. multi-linear regression (see **Methods, Eq 9-12**), which has the advantage that it makes no assumptions about HRF shape^59^. The deconvolution was done over partially overlapping 150-sec windows (75-sec steps; 150 sec typically covered 10 trials) to get adequate temporal resolution for tracking rest states (e.g. the rest epoch in Fig 8a, indicated by the red bar, lasts about 400 sec. Shorter deconvolution windows led to excessive noise). Importantly our design matrix contained not only the spiking regressor but also additional DC and slope terms. The DC term is analogous to the ‘y intercept’ in 1-d linear regression, quantifying an inhomogeneous addition to a homogeneous linear equation. The prediction using the full deconvolved HRF kernels including the DC term matches measured hemodynamics very well overall (**Fig 8a**, compare ‘**Hemo, full pred**’, green with ‘**Hemo, meas**’, black). The phasic portions of the HRFs from deconvolution windows falling in the rest epoch matched each other well and resembled canonical HRFs (**inset ‘HRFs’**, **Fig 8a**; also **Supplementary Fig 5** for an example canonical HRF). They also predicted the high-frequency fluctuations in the measured hemodynamics well (**Fig 8a**, ‘**Hemo, phasic pred**’, red). However, they failed to account for the increase in the mean blood volume, predicting a decrease instead (prediction rising above baseline) which is consistent with the decrease in the local mean spiking. The measured increase in the blood volume was well accounted for, on the other hand, by the additional DC term in the deconvolved kernel (**Fig 8a**, ‘**Hemo, DC pred**’. The slope terms made only small contributions). The same pattern was seen over the full 2.5-hour recording (**Fig 8b**). Linear, phasic predictions from spiking matched the high-frequency fluctuations of hemodynamics during rest epochs, while the deconvolved DC tracked the mean measured hemo (compare **Fig 8b**, ‘DC pred’ with ‘Hemo’ trace in Fig 7a). This result supports the suggestion that the large changes in mean blood volume during rest were likely driven by a mechanism acting in addition to spiking. Similar results were obtained for all extended recording sessions, including ones where the mean spike rate increased during rest.

## Discussion

Our goal is to understand the task-related endogenous component of hemodynamic responses recorded from visual cortex of subjects engaged in cued, predictable tasks. The existence of such responses has been known for more than a decade ^16,33,35–41^, their substantial strength relative to other brain hemodynamic components is well recognized^34^, and recent studies suggest their relevance to sensory processing^33,35^. Yet they have not been adequately studied and little is known about their underlying mechanism or behavioral significance. We earlier reported one particularly prominent task-related response in V1 of alert macaques^31^, which is likely closely related to responses seen in human visual cortex^16,37^. Our work here suggests that this task-related response reflects brain mechanisms of task-engagement. On increasing reward size to get the animal more engaged, the most notable effect, trial by trial, was improved temporal precision: the response became consistently more crisply aligned to task timing. It also became modestly stronger. At a slower timescale, different levels of task engagement led to consistent DC shifts in the mean local blood volume. Notably, high-reward blocks led to consistent decreases in the mean blood volume while low-reward blocks led to corresponding increases. This effect was even more pronounced when the animal disengaged completely from his task and rested with eyes closed. The mean blood volume increased strikingly during these breaks, returning to baseline when the animal resumed working. None of these effects at any time scale could be accounted for by concurrent local spiking. On a methodological note, we recorded the hemodynamic response using intrinsic-signal optical imaging, at a wavelength tuned for measuring cortical blood volume. Such recordings have a much steadier and more reliable baseline than BOLD fMRI, suffering much less from instrumental noise and drift. This allowed us to monitor the response continuously over many hours thus obtaining the reported results, including the prominent DC shifts.

We propose that the task-related hemodynamic response and the effects reported in this paper are linked to brain states of vigilance^55^, in the sense of sustained attention during an extended and possibly repetitive task. This is to be distinguished, on the one hand, from selective visual attention (see the brief review in the Introduction) and, on the other, from non-specific arousal along the sleep–wake axis^55^. Sustained attention is known to fluctuate between states of higher stability which are less prone to error (‘in the zone’), and states that are more unstable and error-prone (out of ‘the zone’; see, e.g., ref^60^). Notably, the state of being ‘in the zone’ is marked by higher regularity and temporal precision: responses ‘in the zone’ show less variability in reaction time, trial by trial, even when the mean reaction time remains unchanged overall ^60^. The state is further enhanced by reward, which leads to even less variability in reaction times, in a manner that appears distinct from increased arousal^61^. This behavioral result may have a physiological analog in our finding of improved temporal precision or regularity of the task-related hemodynamic response during high reward (**Fig 3** and accompanying text). Much remains to be done, to understand the function of the task-related hemodynamic response and its link, if any, to the behavioral correlates of sustained attention. An interesting opening is afforded by the finding that the task-related response may act as a temporally selective attention boost, filtering and binding together disparate sensory features that happen to be presented at the same time as the hemodynamic response^33,62^.

What could be the underlying mechanism? Recall that the trial-by-trial response manifests as a coordinated cycle of constriction and dilation of the arterial system, entrained to task timing^31^. This response could be driven by mechanisms (neuromodulatory? feedback?) with both phasic and tonic components, that could modulate local neural responses while also heightening vascular tone by constricting blood vessels. During a task the phasic constriction could flush out the ‘used’ blood allowing blood vessels to refill. As task engagement weakens, the tonic level of this driver could reduce, relaxing blood vessels passively and increasing the mean cortical blood volume. While there may be many candidate drivers (adrenergic^63^? Astrocytic^64^?) targeting different components of the local vascular network an important potential candidate is dopamine^65^. Visual cortex gets dopaminergic innervation even if it is weak compared to other cortical areas^66,67^, and dopamine is proposed as a mediator of both constriction and dilation of cortical blood vessels through different receptor classes^68,69^. Functionally, dopamine has also been proposed as a mediator of improved perceptual discrimination and response strength in sensory cortex following rewarded trials^70,71^. Exploring these issues through targeted experiments in alert animals would be crucial to understanding brain mechanisms of task engagement and vigilance.

## Methods

### CONTACT FOR RESOURCE SHARING

Requests for resources (e.g. software), data and further information should be sent to Aniruddha Das.

### EXPERIMENTAL MODEL AND SUBJECT DETAILS

Animal use procedures were in accordance with the US National Institutes of Health Guide for the Care and Use of Laboratory Animals and were approved by the Institutional Animal Care and Use Committees of Columbia University and the New York State Psychiatric Institute. Two male rhesus macaques (Macaca mulatta) were used in the study. Access to water was scheduled to training or recording sessions that occurred 3-5 hours per day. Eye fixation and pupil diameter were recorded using an infrared eye tracker (ISCAN^1^).. Before training, each animal was implanted with a stainless steel or titanium head post. After training, craniotomies were performed over the animals’ V1 and glass-windowed stainless steel or titanium recording chambers were implanted for subsequent Intrinsic Signal Optical Imaging (ISOI) in the alert animal (see below). The craniotomy exposed a 20-mm dia area of V1 covering visual eccentricities from ~1 to 10°. The exposed dura was resected and replaced with a soft, clear silicone artificial dura (GE Silicone RTV615 001). Recording chambers and artificial dura were fabricated in our laboratory following published designs^2,3^. Chambers were opened regularly for cleaning and testing for infection and, if necessary, treating, following published protocols^4^.

### METHOD DETAILS

#### Summary

Extracellular electrode recording was carried out simultaneously with intrinsic-signal optical imaging from V1 of alert monkeys performing a periodic visual fixation task. Task and recording methods, described below, are essentially identical to those in earlier papers from our lab^5–8^

#### Task and reward schedules

All experiments were based on a simple fixation task, carried out either under essentially complete darkness or in the presence of visual stimuli. In both conditions, animals held fixation periodically, cued by the color of a fixation spot. (Fixation window: 1.0 to 3.5 deg. diameter, monitor distance: 133 cm; fixation duration: 3–5 sec; trial duration: 9–22 sec; all parameters fixed for a given experiment, but variable between experiments). A juice reward followed every correct (unbroken) fixation, with no time out or other punishment for errors. The primary behavioral manipulation consisted of systematically changing reward size.

For fixation trials in the dark room, the monitor was covered and the fixation point was presented behind a pinhole^8^. Reward sizes were alternated between high (typically 0.45 ml per correct trial, ranging from 0.35 – 0.6 ml) and low (typically 0.15 ml per correct trial, ranging from 0.1 – 0.2 ml. High and low reward sizes were fixed for an experiment; they were selected per day based on the animal’s willingness to work for the low reward). Rewards were alternated in blocks (typically 10 correct trials each; some experiments had longer blocks; some experiments had blocks of variable size. The animal had to correctly complete the full set of trials per block – i.e. not counting error trials – before the reward switched. Trials were grouped into ‘high-reward’ and ‘low-reward’ blocks for analysis. These experiments accounted for the majority of the reported results. For visually stimulated trials (**Fig 5** and **Supplementary Fig 5**)., the animals were passively shown gratings of different contrasts while holding fixation (sine-wave gratings; contrasts doubled in steps ranging typically from 6.25% to 100%; mean luminance = background luminance = 46 cd/m^2^; spatial frequency: 2 cycles/deg; drift speed 4 deg/sec; diameter 2-4 deg; orientation optimized for the electrode recording site. These data are reanalyses of earlier experiments designed to relate hemodynamics to electrophysiology over a wide dynamic range of stimulated responses^5–7^). Reward sizes for these experiments increased progressively from a baseline (typically 0.2 ml per correct trial) to a maximum value (typically 0.6 ml per correct trial) for each successive correct fixation to keep the animals motivated. Again, the lowest reward size per day was chosen based on the animals’ willingness to work. For analysis, trials were grouped into ‘high’ and ‘low’ reward sets relative to the median reward.

#### Intrinsic-Signal Optical imaging (ISOI)

ISOI is based on the finding that in vivo, and in the visible spectrum, changes in light absorption in cortical tissue primarily measure changes in oxy and deoxyhemoglobin in the blood flowing through cortical blood vessels^9–11^. ISOI deduces hemodynamic responses by imaging changes in light reflection at relevant wavelengths off the exposed cortical surface. In effect it is an optical analog of fMRI, albeit restricted to upper layers of exposed cortex. We imaged at 530 nm (green), an isosbestic wavelength that is equally absorbed by oxy and deoxyhemoglobin. Increased absorption of light at this wavelength thus measures increased cortical tissue fraction of hemoglobin, in effect local cortical blood volume, independent of oxygenation state^12^. After the animals had recovered from surgery, we used this technique to image their V1 through the glass-window of the recording chamber, routinely, while they engaged in the fixation task. **Imaging hardware.** Consisted of the following: camera, Dalsa 1M30P (binned to 256 × 256 pixels, 7.5 or 15 frames per second); frame grabber, Optical PCI Bus Digital (Coreco Imaging). **Imaging software:** was developed in our laboratory in C++ based on a previously described system^13^. **Illumination,** high-intensity LEDs (Agilent Technologies, Purdy Technologies) **Lens:** macroscope^14^ of back-to-back camera lenses focused on the cortical surface. Imaging, trial data (trial onset, stimulus onset, identity and duration, etc.) and behavioral data (eye position, pupil size, timing of fixation breaks, fixation acquisitions, trial outcome) were acquired continuously. All data analyses were performed offline using custom software in MATLAB (MathWorks; **RRID:nlx_153890**).

#### Electrophysiology

Electrode recordings were made simultaneously with optical imaging. Recording electrodes (FHC, AlphaOmega; typical impedances ~600–1,000 kΩ) were advanced into the recording chamber through a silicone-covered hole in the external glass window, using a custom-made low-profile microdrive. Recording sites were mostly, but not exclusively, confined to upper layers. Signals were recorded and amplified using a Plexon recording system (**RRID:nif-0000-10382**). The electrode signal was split into spiking (100 Hz to 8 kHz bandpass) and LFP (0.7–170 Hz). Subsequently, an additional analog 2-pole 250 Hz high-pass filter was applied to spiking, effectively eliminating any spectral power overlap between LFP and spiking. No attempt was made at isolating single units and all measured spiking was multiunit activity (MUA) defined as each negative-going crossing of a threshold = ^~^4× the r.m.s. of the baseline obtained while the animal looked at a grey screen. The LFP recording was analyzed to obtain two bandpass-limited measurements, in the beta- and theta-range frequency bands (15-25 Hz, and 3-7 Hz respectively; multi-taper spectral analysis using the Chronux MATLAB toolbox) This gave an LFP measure of (low) vigilance defined by the square root of the ratio of power in theta vs beta.

#### Analysis: pre-processing

The imaging measurement was averaged over the imaged area, frame by frame (frame rate: 7.5 or 15 frames/sec), then divided by the mean value of this quantity for the given experiment (over all trials). This converted the measurement per image frame into 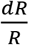 i.e. the fractional change in light reflected off the cortical surface. At the particular imaging wavelength of 530 nm, the negative of this quantity, 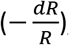, is proportional to the fractional increase in local tissue hemoglobin, i.e. fractional increase in local cortical blood volume^12^. The 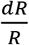 was then detrended and a prominent pulse artifact was filtered out from the measured hemodynamics using Runline (Chronux) with a window of 2 sec. This filtered 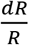 defined the measured hemodynamic response for all calculations.

The pulse artifact was used to estimate the instantaneous heart rate (HR) after upsampling 8x and identifying peak times and thus the local pulse rate. Both the estimated HR and the spiking measurements were then resampled and aligned to the imaging frames. Neither the imaging nor spiking nor estimated HR were further temporally filtered. Importantly, unlike in our earlier papers, we did not high-pass filter to remove slow fluctuations^5–8^, specifically so as to be able to estimate fluctuations over slow time scales of many minutes.

#### Template matching (dark-room experiments: Fig 2)

The amplitude and timing of the task-related response, per trial, were estimated as the height and location of the corresponding peak of a *Template Match*. This *Template Match* consisted of the continuous, normalized dot product of a template with the measured hemodynamic response. The calculation involved the following steps:

- The default template ‘*Tmplt*’ was defined to be the one-trial-long mean hemodynamic recording (z-scored to give ‘*H*(*t*)’) aligned to trial onsets, averaged across all correct trials, and mean-subtracted.
- This *Tmplt* was then slid over *H*(*t*) in unit time steps (at the resolution of the imaging frame rate; e.g. 66.7 ms for 15 frames/s). At every time point *t* the *Template Match* was defined to be the local dot product over the one-trial-long section of *H* centered on *t*, normalized by the (fixed) sum of squares of the *Tmplt* (Fig 2b):

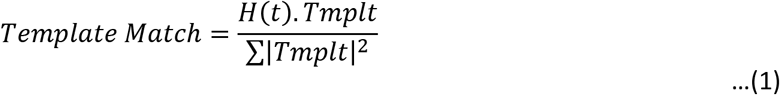 This expression is identical in form to the Pearson’s correlation between the *Tmplt* and the same one-trial-long segment of *H*(*t*), other than the normalization. Thus, like Pearson’s r, this expression would have local maxima where the local one-trial-long segment of *H*(*t*) matched the *Tmplt* in shape (**Fig 2b**). The *Template Match* is also invariant to DC shifs in *H*(*t*) since the *Tmplt* is mean-subtracted and thus integrates to zero over any constant DC. However, unlike Pearson’s r, this expression carries scale information. Pearson’s r has the standard deviation of both arguments in the denominator making it scale-invariant. The *Template Match* on the other hand, with its fixed normalization independent of *H*(*t*), scales linearly with the amplitude of fluctuation in *H*(*t*). Thus peaks of the *Template Match* carry information about both timing and amplitude of the task-related response per trial.
- For computational efficiency in Matlab the above expression was rewritten as the normalized convolution of *H*(*t*) with the time-reversed version of the template *Tmplt*:

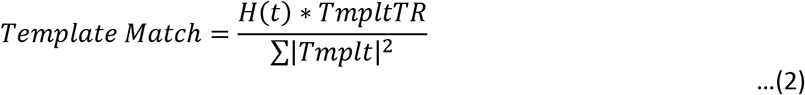

where *TmpltTR* is the time-reversed *Tmplt*, i.e. *TmpltTR* = *Tmplt*(−*t*) and the symbol ‘*’ denotes convolution. The denominator for normalization remains unchanged. This expression translated to the following script using Matlab functions *conv* and *sum*:

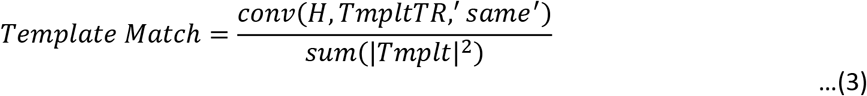 Peaks of *Template Match* were then identified as zero crossings of the first derivative at points where the second derivative was negative (marked by red dots in Fig **2b**).
- Alternate template matches used the same formalism but with different definitions of *Tmplt*. Thus the *Tmplt* in **Supplementary Figure 2** was defined as the one-trial-long mean *H*(*t*) aligned to a point one quarter trial ahead of trial onsets, averaged across all correct trials, and mean-subtracted. All other steps were the same

#### Template matching (with visual stimuli present: Fig 5, Supplementary Fig 5)

This involved two separate sets of steps.

1. Estimating the task-related response from the net recorded hemodynamic response
  - We modeled the net recorded response as a linear sum of stimulus-evoked and task-related components. The stimulus-evoked response was modeled as the convolution of concurrent spiking with an *HRF* kernel. The task-related component was estimated iteratively. Our earlier approach^6^ had modeled it as a stereotyped task-related function (‘*TRF*’) that was identical in timing and amplitude for each correct trial. Here, however, we specifically need to estimate trial-by-trial variations in response timing and amplitude. As a first step, we assumed that the *TRF* had a fixed shape that could be estimated from the mean across trials. Optimal mean *HRF* and *TRF* kernels were obtained by fitting the mean recorded responses, separated by contrast, with the following equation, identical to Eq 1 in ref^6^:

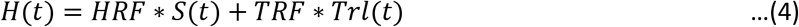 *H*(*t*) is the recorded hemodynamics and *S*(*t*) the concurrently measured spiking, and *HRF* * *S*(*t*) models the stimulus-evoked response. The second term on the RHS models the task-related response as a *TRF* kernel convolved with the set of delta functions at trial onsets, *‘Trl*(*t*)*’*. The symbol ‘*’ denotes convolution over time. The *HRF* kernel was parametrized, as before^5,7,8^, as a Gamma-variate function of time *t*:

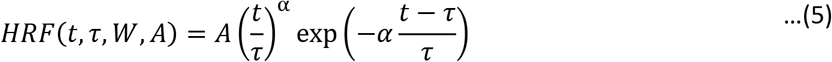 The *HRF* parameters fitted during optimization are the amplitude *A*, time to peak *τ*, and full width at half maximum *W* ^8,15,16^. The factor 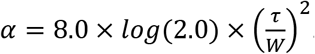. The *TRF* kernel was parametrized as the finite sum of a Fourier time series:

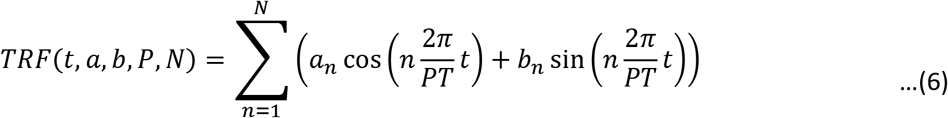 Although the Fourier series was based on the trial period *T*, the fundamental Fourier period was allowed to vary as a fraction *P* of the trial period and optimized in the fit. The parameters *a_n_* and *b_n_*, (with *n* ranging from 1 to the total number of terms in the Fourier series, *N*) are the pairs of cosine and sine coefficients, respectively, for the *n*th Fourier term. We showed earlier that only the fundamental and first harmonic, i.e. *N* =2, carry significant information^6^. Thus there are 8 parameters in the model: 3 for the *HRF*, the 2 pairs of *a_n_* and *b_n_*, and *P*.
  - All parameters were optimized simultaneously by matching the predicted to the measured hemodynamics using a downhill simplex algorithm (*fminsearch*, MATLAB methods as in ref^7^). To keep contrast information, we made concatenated sequences of the mean response per distinct contrast, randomized per contrast (same random sequence for hemo, and spiking), and over multiple blocks (an arbitrarily large number 52, about 100x larger than a single *HRF* kernel convolution length, to minimize edge effects; we only matched traces two convolution lengths in from the edge). The error to be minimized was defined as the normalized sum squared error 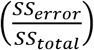 calculated separately per contrast and then averaged over all contrasts including the blank. This was intended to give equal weight to the fractional error at each stimulus contrast. The goodness of fit *R^2^* for the optimal prediction was defined as the coefficient of determination 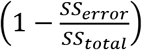 calculated separately per contrast and then averaged across contrasts. This, again, was intended to give equal weight to errors at each contrast. In order to reduce the chances of getting caught in a local minimum we started with large sets of initial parameter values, independently covering an order of magnitude for each fitted parameter. The fits were robust and converged to the same optimal parameters from multiple starting values, giving us confidence that we had reached global and not local minima.
  - Next we used the optimal fitted *HRF* thus obtained from the mean hemo and spiking, averaged per contrast, to get a continuous estimate of the exogenous, stimulus-evoked component of measured hemodynamics. This was done by convolving the optimal *HRF* with the measured spiking:

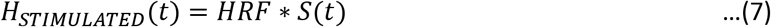
  - This estimate of *H_STIMULATED_* was subtracted from the full measured hemo to get an estimate of the endogenous, task-related component of hemodynamics as the residual not accounted for by spiking:

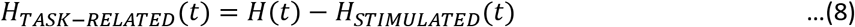
2. Estimating task-related response peaks and amplitudes.
  - The estimate of the task-related response *H_TASK_*_–*RELATED*_(*t*) as defined above was then used, exactly like the full hemodynamic response in the dark-room, to estimate response times and amplitudes trial by trial. As template, we used the optimal *TRF* obtained above by fitting to mean responses. The steps for template matching, and identifying and analyzing peaks were identical to those outlined in Equations 1-3, with the *H*(*t*) being replaced with *H_TASK_*_–*RELATED*_(*t*) and the *Tmplt*(*t*) being replaced with the optimal *TRF*. Peak times and amplitudes, per trial, were obtained exactly as in the dark-room template match.

#### Tracking measured variables as animal switches from alert and task-engaged to disengaged with eyes closed (Fig 7)

To track slow changes in all measured variables, we downsampled the data. Data were averaged using a 15 sec box car that corresponded roughly to a single trial, then decimated 4x giving, in effect, a smoothed 60-sec sample rate. Along with MUA spike rates at individual electrodes, and the measured hemodynamics, the following measurements were thus tracked:

1. ‘Eyes closed’: Fraction of time over the 60-sec averaging window that eyes are closed. Eye closure was monitored using the output from the IR eye tracker. All blinks or eye closures appeared as sequences of missing points or ‘rails’ (saturated output). Spontaneous eye blinks in macaques last roughly 200 ms (see ref^17,18^). Our own data showed a bimodal distribution with blink durations peaking either at 200 ms or multiple seconds to minutes. Thus sequences of missing points lasting < 500 ms were considered regular spontaneous blinks during the alert state and were marked as having duration = 0. Sequences lasting > 500 ms were categorized as eye closures and their durations were included in the moving average.
2. ‘LFP measure’: square root of the ratio of spectral band – limited power in the theta (3-7 Hz) and beta (15-25 Hz) frequency bands, each normalized by its standard deviation over the entire experiment. We chose this particular ratio to get a measure that was high during epochs of low alertness to match ‘Eyes closed’ since theta power increases sharply on transitions from high to low alertness or to sleep^19^. We took the square root of the ratio to compress the measure to ~0-1, to make it comparable to ‘Eyes closed’. There was no attempt to separate the resting state more finely into sleep stages since the goal was a broad separation into states of ‘alert and task-engaged’ vs. ‘resting’ with a time resolution of 60 sec.
3. ‘Spk1-Spk2 Corr’: Pairwise correlation of MUA spike rate from individual electrodes. Calculated over a one-second moving window.
4. ‘HR’: Heart rate: Obtained from the pulse artifact in the measured hemodynamics, after upsampling 8x and identifying peak times and thus the local pulse rate. This instantaneous pulse rate, estimated at pulse time points, was then interpolated with spline smoothing to the imaging time base (7.5 Hz or 15 Hz sample rate depending on the experiment)

#### Deconvolution, i.e. multilinear regression (Fig 8)

We started with the assumption that the measured hemodynamics *H*(*t*) can be predicted from local spiking *S*(*t*) using a homogeneous linear equation, along with two inhomogeneous terms: a DC offset, and a linear slope (in the vector space of each 150-sec window, at the resolution of the camera frame rate):

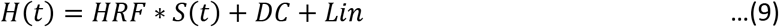

There are no assumptions about the shape of the *HRF* other than that it does not extend more than 10 sec prior to time 0, and is back to baseline about 25 sec after time 0. Using the formalism of deconvolution, this expression can be rewritten as a matrix equation

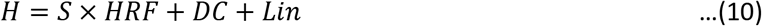

Where *H* is a column vector of recorded hemodynamic responses (at the temporal resolution of the camera frame rate, 15 Hz), *S* is the spiking regressor expanded into a stimulus convolution matrix (SCM) ^20,21^, the symbol ‘×’ indicates matrix multiplication and *HRF*, *DC* and *Lin* here refer to the same terms as in Eq 10 but expressed as column vectors. The SCM was constructed as a Toeplitz matrix comprising a horizontal concatenation of spiking column vectors, with circular time shifts ranging from −10 sec to +25 sec relative to t=0. Formally, the SCM *S* can be extended (‘*S_e_*’) to incorporate the *DC* and *Lin* by horizontally concatenating the two additional column vectors: a column of ones for the DC, a linear ramp from −1 to 1 for the slope. The corresponding *HRF* can be formally extended (‘*HRF_e_*’) by two coefficients, one for the *DC* and another for the *Lin*.

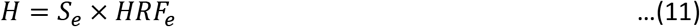

Assuming that any noise is Gaussian and zero-mean, the optimal deconvolved *HRF_e_* can then be estimated using a least-squares solution to the linear regression^20,21^:

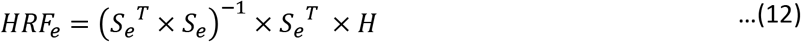

Where the superscript ‘^*T*^’ indicates the matrix transpose, ‘^−1^’ indicates the matrix inverse and ‘×’ indicates matrix multiplication. The full prediction using the optimal deconvolved *HRF_e_* kernel is then computed using Eq 11. Similarly, just the (linear, homogeneous) phasic prediction from spiking is obtained by taking the matrix multiplication over all column vectors of *S_e_* save the last two, with all coefficients of the deconvolved *HRF_e_* save the last two. Conversely, just the DC term or just the slope term are obtained by appropriately multiplying the last two column vectors of *S_e_* with the corresponding last two coefficients of the deconvolved *HRF_e_*.

#### Getting bootstrap estimates for significance (p values)

All comparisons between distributions of amplitudes or peak times were tested for significance by bootstrapping, typically using 10,000 resamples with replacement. In cases with different numbers of trials for high and low reward, the smaller number of trials was chosen to make the bootstrap comparison. For comparing response amplitudes (e.g. **Fig 3b**) we tested for the median of high-reward amplitudes being > that for low-reward amplitudes over the set of all resamples, against the null hypothesis that this difference has zero mean. We also tested for the complement, i.e. that median of low-reward amplitudes > that of high-reward amplitudes. For comparing widths of peak time distributions (e.g. **Fig 3a**) we first calculated the 2σ width (specifically, the +/- 34^th^ percentile around the median, given the non-normal distribution) of each bootstrapped set of peak times, separately for high and low reward. We then tested for 2σ for high reward < for low reward over the set of all resamples, against the null hypothesis that the difference has zero mean. We also tested for the complement, i.e. that 2σ for low reward < for high reward.

#### Fitting spiking to dark-room hemodynamic response using gamma-variate *HRF* (Supplementary Fig 1)

To link to spiking, the dark-room response was modeled as a homogeneous prediction from spiking, fitted by optimizing a gamma-variate *HRF* kernel using *fminsearch* as described above:

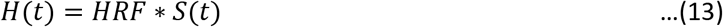

The fitting was done separately for the high-reward and low-reward trials at each recording site. Stimulus-evoked responses were fitted using a model with a task-related component: Eq 4-6. In each case the optimal fitted *HRF* kernel was then convolved with the continuous recorded spiking response to give a continuous prediction. Since the spiking response included both high-reward and low-reward segments, the prediction included sections of ‘same’ prediction (e.g. low-reward spiking convolved with the low-reward kernel) and sections of cross-predictions (e.g. high-reward spiking convolved with the low-reward kernel).

---

**Supplementary Figure 1.**
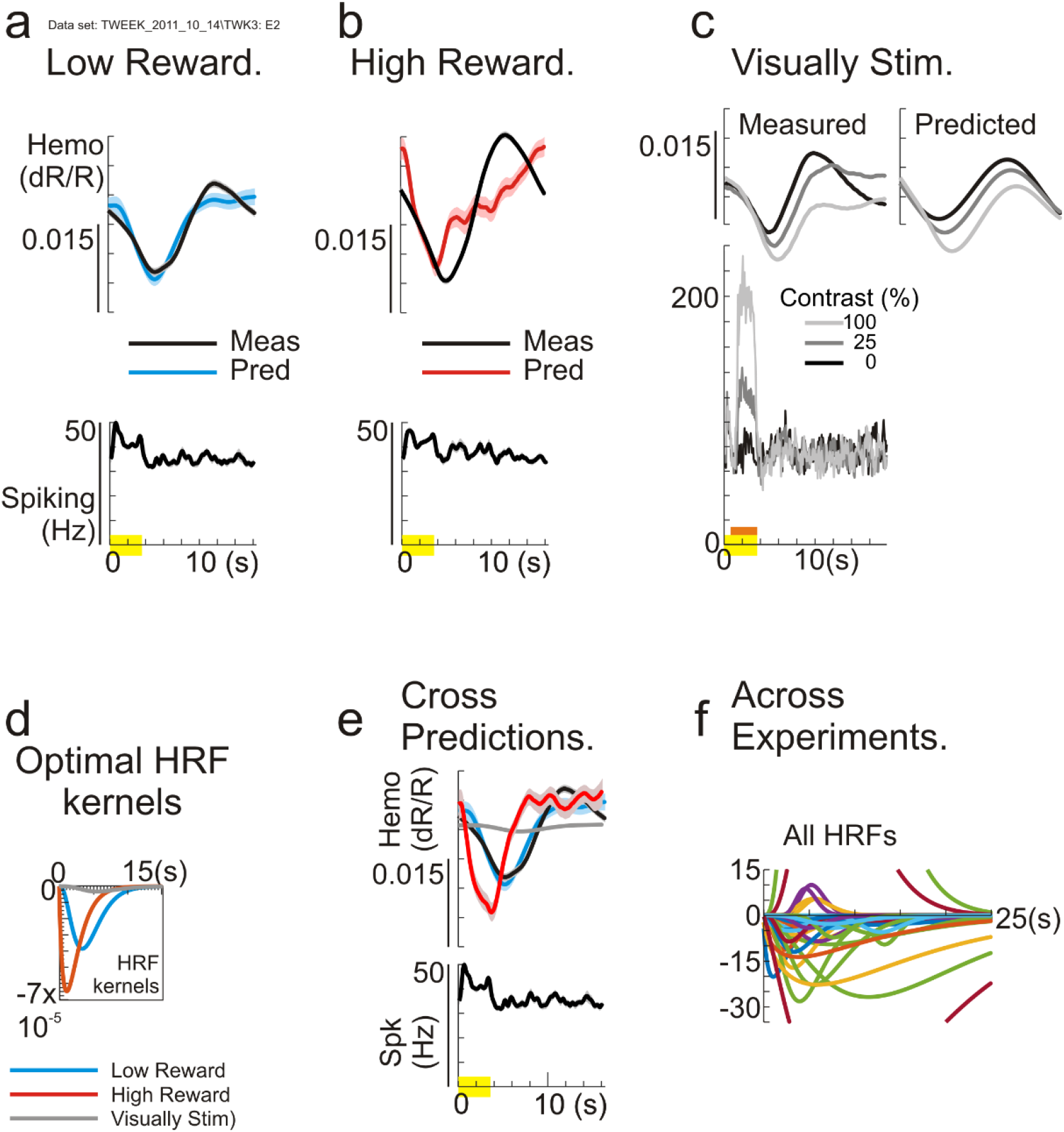
Local spiking, while appearing to predict mean hemodynamic responses in individual recording conditions, is a poor and unreliable predictor of task-related responses overall. **a,b:** Mean measured responses and optimal predictions for low-reward and high-reward trials, respectively, of a data set recorded in the dark room task. In each case the lower panel shows the mean measured spiking while the upper panel shows the mean measured hemodynamics as well as the prediction from spiking using the corresponding optimal fitted gamma-variate HRF kernel (see color code in each column). Low reward: N=148 trials, R^2^=0.73 for the optimal prediction. High reward: N=140 trials, R^2^=0.42. **c:** A separate set of visually stimulated trials at the same recording site, using visual stimuli consisting of optimally oriented drifting gratings at different contrasts, as indicated by the grayscale coding (orange bar below depicts the visual stimulation period). Again, the top panel shows mean measured hemodynamics and optimal predictions grouped by stimulus contrast; predictions are shifted to the right for visibility. N=141 trials total, i.e. 47 trials / contrast. R^2^=0.95. **d:** The optimal fitted gamma-variate HRF kernels for the three recording conditions, color coded as shown. Note how poorly they match each other. **e:** Comparing the measured low-reward hemodynamics to predictions using the low-reward dark-room set of spiking responses (as in Panel a), but with different optimal HRF kernels; from low-reward, high-reward and stimulus-evoked sets. The cross predictions are poor (R^2^ of prediction using high-reward HRF=- 0.014; stimulated HRF =-0.011). **f:** Optimal HRFs from the full set of dark-room experiments, normalized in each case to the amplitude of the corresponding visually stimulated HRF. (N=56: pairs of high and low reward HRFs for each of 28 sets with electrode recordings). Scale truncates some HRFs of high absolute amplitude to help visualize those of smaller amplitude. Colors are arbitrary (Matlab default). The different optimal HRFs match each other poorly, with some even reversed in sign. This makes cross prediction meaningless and suggests that apparently good predictions of mean responses in individual experiments are fortuitous.

**Supplementary Figure2:**
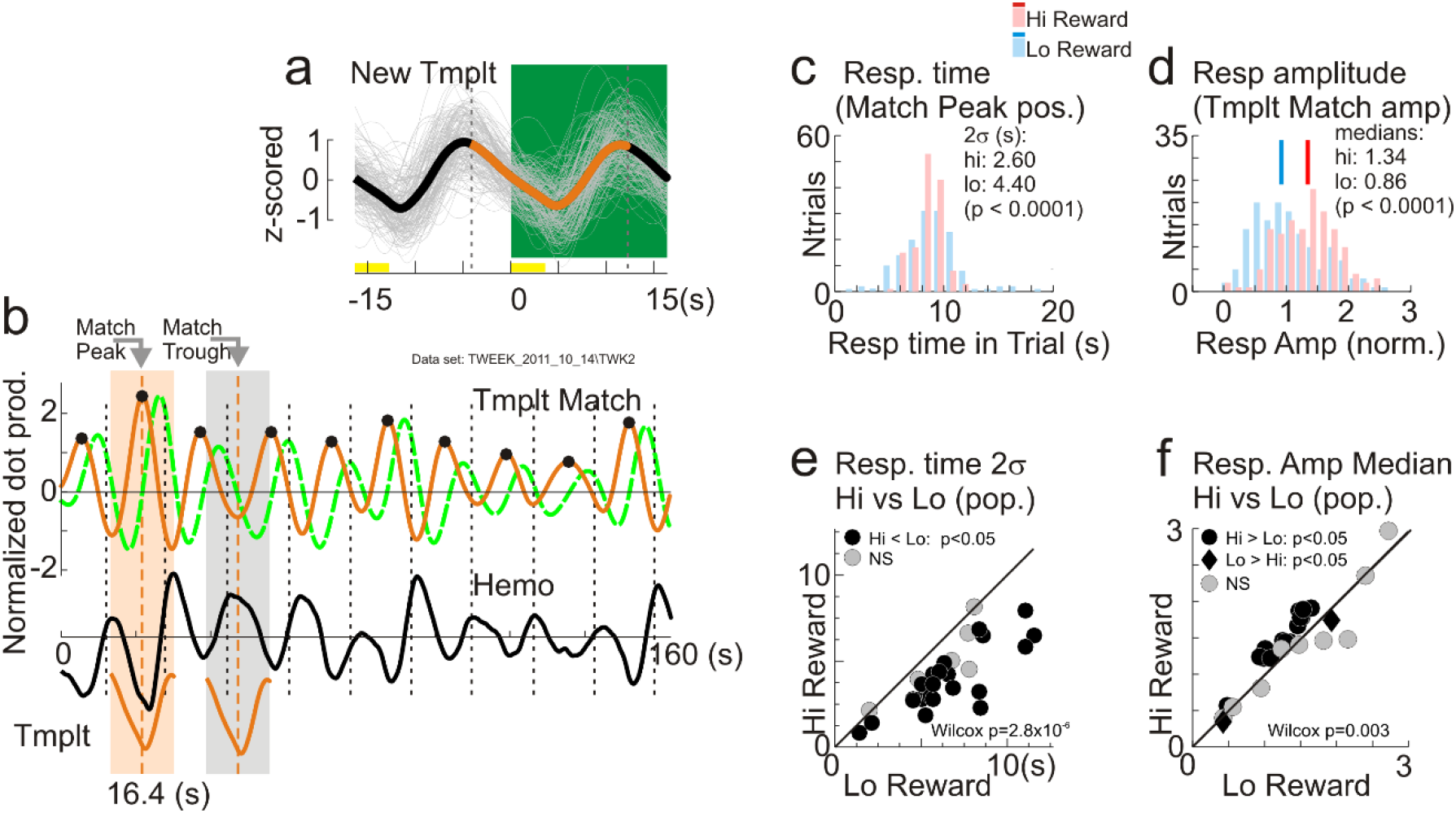
Increase in temporal precision with reward size is not sensitive to the choice of template used to estimate response time and amplitude **a-d:** Same example dataset as in Figs 2, 3. **a:** Orange, alternate template defined as the mean hemodynamic response across correct trials, aligned to a time point one quarter cycle ahead of trial onset (i.e. starting at the dashed vertical line 4.1 sec ahead of time 0. Single trials shown in gray). Green background (time points 0 – 16.4 sec) marks the timing of the earlier template, for comparison (see Fig 2b, ‘Tmplt’). **b:** New template match (orange, ‘Tmplt Match’, upper row), illustrated using the same segment of recorded hemodynamics (‘Hemo’) as in Fig 2b. The earlier template match from Fig 2b is shown alongside for comparison (green, dashed line). Black dots identify the peaks of the new Template Match, marking locations where the Hemo is locally best phase-matched to the new template (see ‘Match Peak’, compared to ‘Match Trough’). **c:** Distributions of response times, defined as the positions of the new template match peaks. Compare with Fig 3a. **d:** Distributions of response amplitudes using the new template match. Compare with Fig 3b. **e, f:** New response timing distribution 2σ widths, and amplitude medians for high vs. low reward trials across all experiments, including p values from Wilcoxon signed rank test for the pairwise comparisons. Compare with Figs 3c,d

**Supplementary Figure 3:**
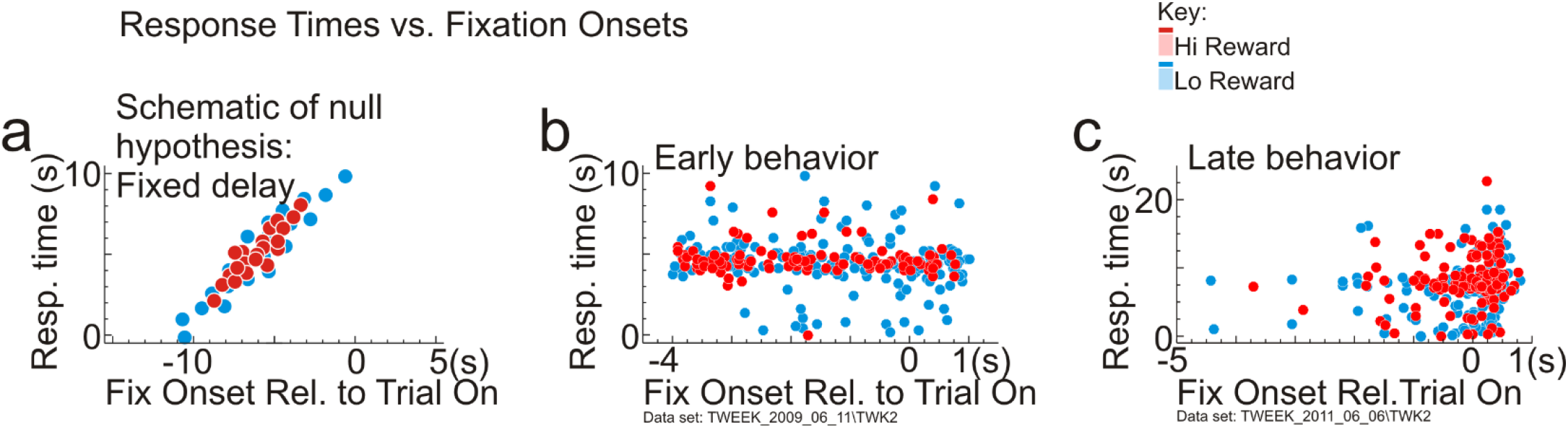
Response timing does not correlate with fixation onset. **a:** Simulation of the null hypothesis. The task-related response has a stereotyped time course following the onset of fixation. Response times would then have a constant delay following fix onset, leading to a linear relation between the two with unity slope (the delay was taken to be 10 sec for this simulation). The observed tighter clustering of response times for high reward could result from a corresponding clustering of fixation onsets (consider projection of red dots, vs. blue dots on the Response Time axis). **b:** Relationship between measured response time, estimated as usual with a template match, and fixation onset, in an early recording session. Animals tended to hold fixation for extended periods prior to trial onset, even across multiple trials. **c:** Relationship between response time and fix onset in a late recording session. Animals tended to move their eyes a lot during intertrial intervals, fixating shortly before trial onset. For both cases ‘b’ and ‘c’ response times were independent of fixation onset and very different from the pattern expected for the null hypothesis. In both data sets response times for high-reward trials showed visibly lower scatter independent of fix onset.

**Supplementary Figure 4:**
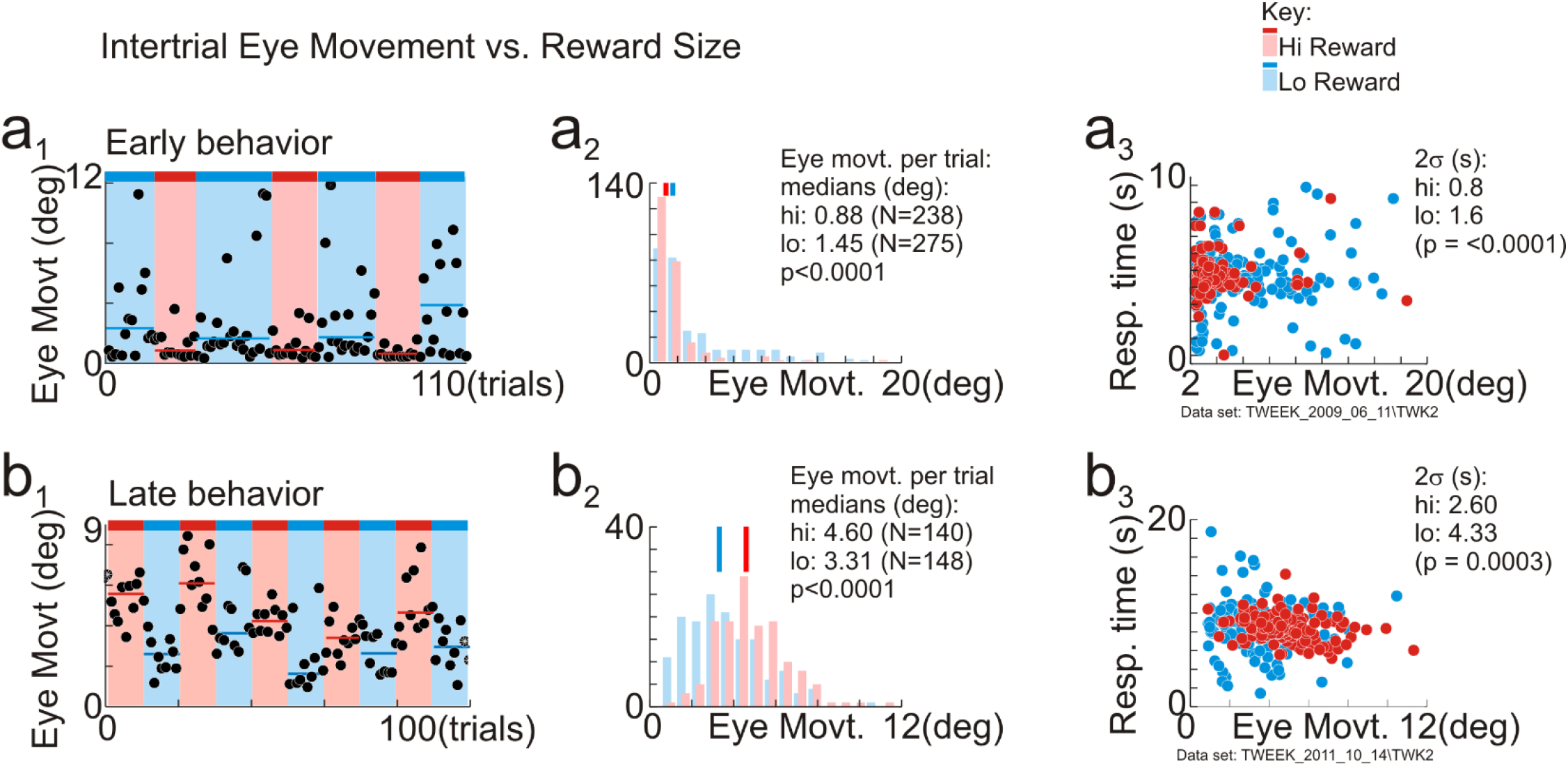
High reward does not correlate with tighter eye movements. **a1**: Mean radial eye movement per intertrial interval in an early recording session. Each dot represents a single trial (mean eye movement during 7-sec intertrial intervals per 9 sec trial.). Horizontal lines, median eye movement per block of high or low reward (blocks with varying numbers (13-31) of correct trials each). Intertrial eye movements were higher in low-reward blocks. **a2:** Histogram of mean eye movement per trial. **a3:** Relationship between response time and eye movement per trial, colored by reward size. **b1:** Mean radial eye movement in a later recording session (12-sec intertrial intervals in 16-sec trials; alternating blocks of 10 correct trials each; all other conventions as in panel a1). Eye movements were higher in high-reward blocks. **b2:** Corresponding histogram of mean eye movements per trial. **b3**: Response time vs eye movement, per trial colored by reward size. Low reward leads to wider scatter of response times in both panels (a3) and (b3) despite opposite effects on intertrial eye movement.

**Supplementary Figure 5:**
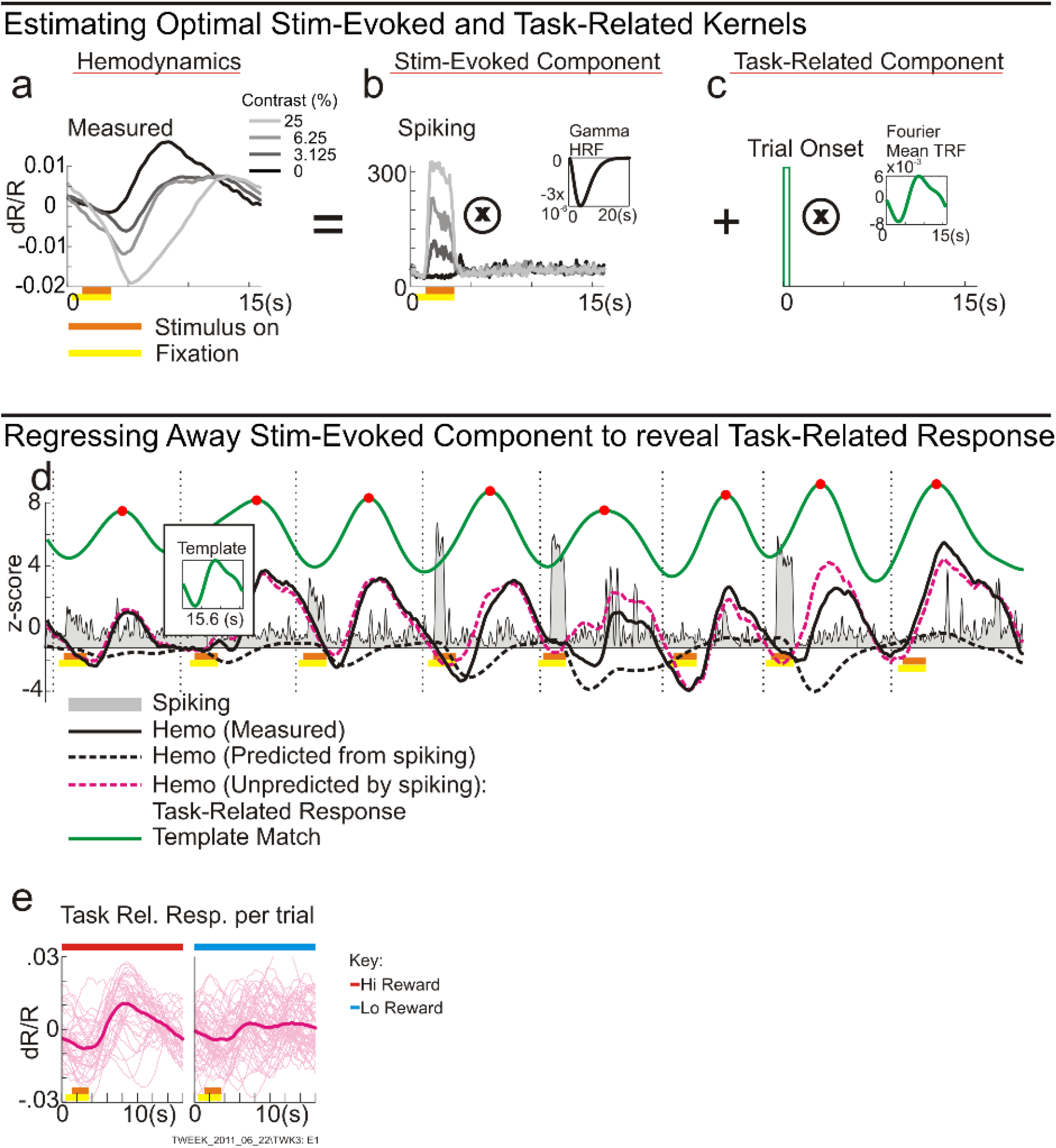
Estimating Task-Related Response and its Template Match in the Presence of Visual Stimulation (one example data set). **a-c:** Estimating optimal fitted parameters (see **Methods, Eq 4-6**). **a:** The mean hemodynamic response per stimulus contrast (see key), averaged across trials. The response is modeled as the sum of: the stimulus-evoked component (b), and the task-related component (c). The stimulus-evoked component is modeled as the convolution 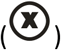 of the measured spiking with a gamma-variate HRF kernel (inset). The mean task-related component is modeled as the convolution of delta functions at trial onset with a ‘Mean TRF’ kernel comprising a partial Fourier sum with its fundamental at the trial period (inset). Earlier work showed that the fundamental and the first harmonic terms of the Fourier series are adequate. Insets show the optimal fitted Gamma Variate HRF (in b) and optimal mean TRF (in c) respectively. **d:** Set of traces illustrating the process of estimating the residual task-related response, and then estimating its timing and amplitude per trial by matching to a template (see Methods, **Eq 7-8**). ‘Spiking’, ‘Hemo’: full measured responses, individually z-scored. ‘Hemo (predicted from spiking):’ is the convolution of the spiking response with the optimal fitted HRF ((b), inset). Subtracting this from the measured Hemo gives the residual ‘Hemo (Unpredicted by spiking)‘ which we defined to be the task-related response. The moving-window dot product of this residual with the template (the optimal fitted mean TRF ((c), inset) gives the ‘Template Match’ (shifted up for visibility). Timing and amplitude of task-related responses, per trial, are defined to be the location and height of each Template Match peak, as for the dark-room task. **e:** Set of all residual task-related responses, converted back from z-scored values, separated into trials grouped by reward size. The same data are shown in Fig 5a.

**Supplementary Figure 6:**
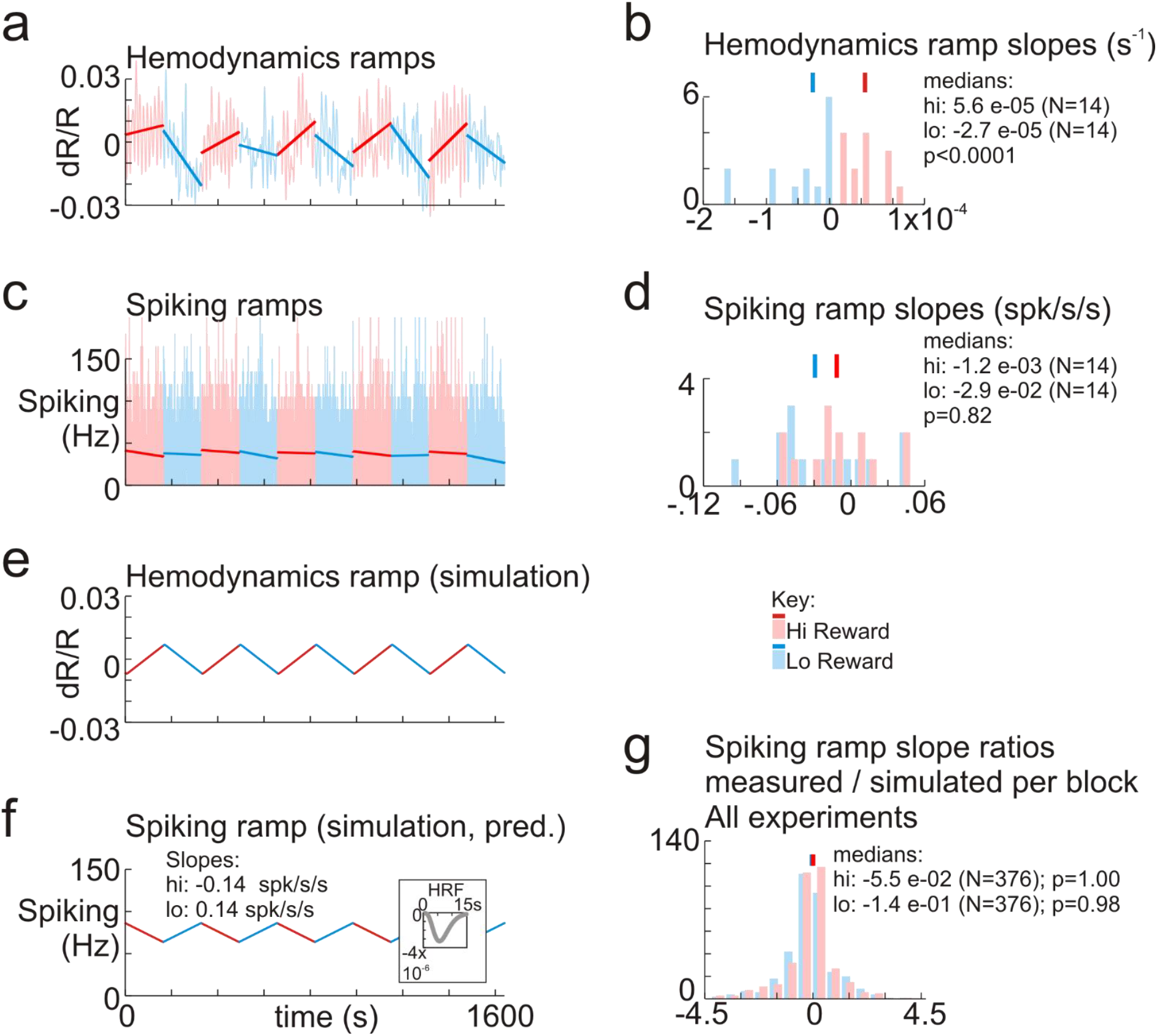
Ramp-like drifts in local blood volume are not accounted for by slow changes in local spiking**. a, c:** Hemodynamics and spiking, respectively, showing correct trials from alternating blocks of high and low reward. Lines show regression fits per block. Same data set as Figs 2,3. **b, d**: Histograms with slopes of regression fits from (a), (c). **e:** Simplified simulation of slow mean hemodynamic responses: triangle wave of matching period, with slopes equal to the median (absolute) slopes of the regression lines in (a) (=4.1×10^-5^/sec). **f:** Simulated spiking response that generates the model hemodynamic response in (e) on convolving with the visually stimulated HRF for this recording site (see ‘HRF kernels, Supplementary Fig 1; also, **Methods**). Measured spiking regression slopes (d) are only ~ 4x weaker than those in the simulation; but they do not alternate in sign with reward size. **g:** Distributions of the ratios of measured spiking regression slope / block, to the slope of the corresponding simulation, as in (e), (f) across all experiments (N=752 blocks of 10 trials each, 376 blocks/reward size. From N=11 experiments with electrode recordings and at least 10 blocks per reward size. p values test for the probability of the distributions being centered on zero (bootstrap, 10000 resamples)

Contributions
MMBC: Experimental design, recordings, data analysis, initial analysis of slow responses
BL: Experimental design, recordings, data analysis
YBS: Initial finding of response variation linked to reward, and rest vs. alert engagement
AD: Overall project design, writing manuscript.

